# Targeted gene transfer into developmentally defined cell populations of the primate brain

**DOI:** 10.1101/2025.04.11.648413

**Authors:** Ana Rita Ribeiro Gomes, Natalie Hamel, Surjeet Mastwal, Naim Wright, David C. Ide, Christopher T. Richie, Ted B. Usdin, Kuan Hong Wang, David A. Leopold

## Abstract

The primate brain possesses unique physiological and developmental features whose systematic investigation is hampered by a paucity of transgenic germline models and tools. Here, we present a minimally invasive method to introduce transgenes widely across the primate cerebral cortex using ultrasound-guided fetal intracerebroventricular viral injections (FIVI). This technique enables rapid-onset and long-lasting transgene expression following the delivery of recombinant adeno-associated viruses (rAAVs). By adjusting the gestational timing of injections, viral serotypes, and transcriptional regulatory elements, rAAV FIVI allows for systematic targeting of specific cell populations. We demonstrate the versatility of this method through restricted laminar expression in the cortex, Cre-dependent targeting of neurons, CRISPR-based gene editing, and labeling of peripheral somatosensory and retinal pathways. By mimicking key desirable features of germline transgenic models, this efficient and targeted method for gene transfer into the fetal primate brain opens new avenues for experimental and translational neuroscience across the lifespan.

## Introduction

The abundant use of genetically modified mouse lines has transformed the study of brain anatomy and function by enabling the systematic examination of molecules or circuit elements under genetic control^1–4^. Examples of analogous germline modification of rats^5,6^, ferrets^7,8^ and nonhuman primates^9–14^ have begun to broaden the repertoire of mammalian experimental and disease models. Among these, developing nonhuman primate genetic models is of particular importance for understanding primate-specific features and disorders of the human brain^15^. However, the establishment of such models faces significant hurdles^16,17^. For one, generating transgenic primates is costly and labor intensive, requiring maintenance of sufficiently large colonies for efficient breeding. Furthermore, while molecular tools and genetic constructs are advancing rapidly, the long gestational and prepubertal periods of primates requires many years of effort to establish a given genetic line. This temporal mismatch has resulted in most emphasis being placed on developing transgenic primate disease models^14,17,18^, with fewer resources devoted to creating genetic lines for basic neuroscience research. As a result, research areas such as primate neurodevelopment, which stand to benefit from such tools and are highly relevant to human disease, remain largely unexplored. Thus, there is a need to consider complementary approaches to germline modification to harness the power of genetic methods in nonhuman primates for basic and translational neuroscience.

An important alternative to germline modification is the somatic introduction of transgenes into a living organism. In experimental neuroscience, viral tools have been routinely used for gene delivery, either by inserting genetic material into the host chromosome or by creating persistent DNA concatemers in the nucleus (episomes) that support transcription^19^. For systems neuroscience, transduction from most *in vivo* viral delivery methods is localized, confined to tissue immediately surrounding an injection site. However, recent advances in viral vector administration have enabled broader coverage across the central nervous system^20^. Notably, delivering recombinant adeno-associated viruses (rAAVs) through the cerebrospinal fluid (CSF) increases the breadth of transduction, particularly when the vector is infused in developing animals. Commonly used in mice, rAAV delivery into the CSF is most effective for achieving broad transgene expression within a brief postnatal window of a few days, after which transduction becomes less efficient and more localized^21^. Given the experimental and biomedical opportunities potentially afforded by widespread gene transfer in the primate brain, we sought to translate this cerebroventricular injection approach to the marmoset monkey. Because the corresponding developmental stages in marmosets and other primates occur *in utero*^22^, we designed a method to achieve gene transfer before birth, using ultrasound guided targeting to inject viral particles into the cerebroventricular system of fetal brains.

The present study describes a minimally invasive fetal intracerebroventricular viral injection (FIVI) procedure, initially developed in rats (*Rattus norvegicus*) and adapted to marmosets (*Callithrix jacchus*), to deliver rAAV at early stages of brain development to achieve broad gene transfer across the primate cerebral cortex. We show that a single injection into the fetal CSF results in rapid, robust, widespread, and enduring transgene expression throughout the brain. By varying the gestational timing of injections and the rAAV serotypes, it is possible to systematically target specific cell populations, thus opening the door to new modes of circuit-based study of the primate brain. We further describe experimental results highlighting several areas of opportunity, including the use of FIVI for intersectional methods using Cre recombinase, CRISPR-based gene editing, and the capacity for gene transfer into sensory pathways outside the brain. We conclude that the FIVI method is straightforward and versatile, offering new opportunities to label developmentally defined cell populations in nonhuman primates for a broad range of experimental topics for systems and translational neuroscience research.

## Results

### Ultrasound-guided viral injections into the fetal brain

To efficiently inject material into the fetal brain, we developed an adaptable system to hold the pregnant female marmoset or rat, an ultrasound (US) transducer, and a penetrating transcutaneous guide tube (**Fig. 1a**). This apparatus was designed with multiple degrees of adjustment to accommodate the variable orientation of the fetal cranium in the uterus, and to allow real-time US visual feedback and realignment of the injection path.

**Fig. 1:**
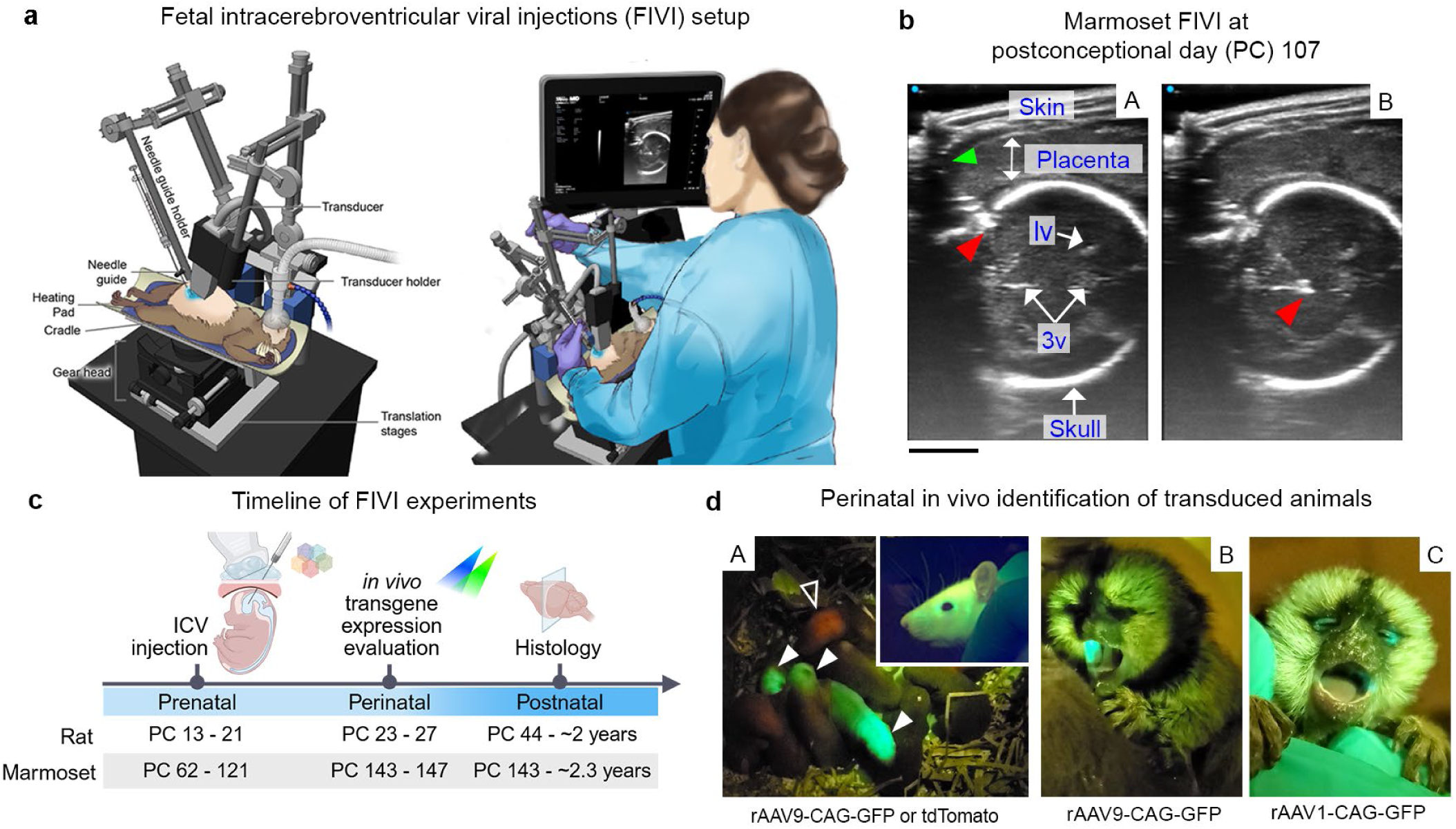
Fetal transduction through ultrasound-guided intracerebroventricular viral injection (FIVI). **a**, Schematic of the FIVI experimental setup and injection apparatus, highlighting its main components, designed for both rats and marmosets. The ultrasound machine, geared head and translational stages allow real-time visualization for precise fetal orientation and injection (see Online Methods for details). Illustration by NIH Medical Arts. **b**, Using high frequency ultrasound linear transducers, the needle can be visualized advancing through a marmoset fetal skull at PC107 (A) and into the brain (B), with delivery of the viral particles into the midline ventricles (see also Supplementary Video 3). The placement level of the guide tip, which is not visible in the panel due to low echogenicity, is marked by a green arrow, while the needle tip is indicated by a red arrowhead. lv: lateral ventricle; 3v: 3^rd^ ventricle. Scale bar, 5 mm. **c**, Timeline of experiments in rats and marmosets. FIVI was successfully applied at various stages, accommodating skull thickening and ventricle volume variations. ICV: intracerebroventricular; PC: postconceptional day. Figure panel created in BioRender. **d**, Reporter expression was visualized in vivo during the perinatal period. In rats (A), reporter gene expression (GFP, closed arrowhead; tdTomato, open arrowhead) is visible in the nervous system through the skull and skin after PC19 injections, with additional expression in muscle. As fur grows and skin thickens, visibility decreases under a flashlight, with expression most prominent in fur-free or thin-skinned areas like around the eyes and snout, as seen in the inset of a PC55 rat injected at PC19. In neonatal marmosets (postnatal day 0), following a PC96 (A) and PC116 (B) injections strong reporter expression can be visualized inside of the mouth and around the eyes. Fainter expression can also be observed in the body (e.g., neck, not shown), though visualization is more challenging due to the darker skin and fur present at birth and varies depending on the injection date and serotype combination.

Detailed step-by-step instructions on assembling the setup and on the procedure, including animal preparation and anesthesia, are given in Online Methods. Briefly, following isoflurane anesthesia induction, the pregnant female is positioned in the cradle in a supine position (**Fig. 1a**). An aseptic environment is created over the shaved abdominal surface, which is covered with sterile US gel. Prior to positioning the transcutaneous guide tube, the experimenter initially determines the location and orientation of the fetus by adjusting the US transducer and observing the corresponding images on the display. The guide tube is supported over the abdomen by a custom micromanipulator arm. It is angled within the US plane, just underneath the edge of the transducer, which is supported by a separate micromanipulator arm. This positioning allows for visualization of the guide tube within the imaging plane, facilitating precise alignment with the fetal target. The guide tube should be visible as it is advanced slowly through the gel and pushed gently against the skin. The guide tube is also electrically insulated along its length, leaving a small area at the tip through which current can be directed using an electrocautery device. To penetrate the skin, a few briefly applied current pulses are delivered through the guide tube tip, which can then be advanced smoothly through the thick abdominal wall without displacement of the fetus (see Online Methods and Supplementary Vids. 1 and 2 for details). This straightforward step greatly simplifies and shortens the FIVI procedure, eliminating the need for skin incision or surgical exposure.

With the transcutaneous guide tube held firmly in position, the injection needle is introduced to the target under ultrasound guidance (**Fig. 1b**). First, the Hamilton syringe needle is manually advanced through the guide tube, abdominal muscle, and uterine wall into the uterine lumen adjacent to the fetal skull. At this stage, small micromanipulator adjustments are sometimes needed to optimize the positioning of the transducer, guide tube, or animal for the most effective targeting of the cerebral ventricles. Once optimized, the needle is manually advanced through the cranial wall toward the ventricular target. We found that this step is best achieved by initially applying a small amplitude mechanical jolt that displaces the needle tip 1-3 mm into the intracranial space and subsequently advancing the needle into the ventricle (Supplementary Vid. 3). Following this ultrasound-guided positioning of the needle tip, the viral particles are infused over 1-2 minutes. The needle is left in place for 2-3 minutes to allow fluid dispersion before retraction. In general, we injected volumes of 5-10 µL in the rat and 10-60 µL in the marmoset over the course of approximately 20-60 seconds. For both rats (typically targeting 6-9 fetuses) and marmosets (typically targeting 2-3 fetuses), the multiple injection procedure usually required between 1.5 and 2 hours of isoflurane anesthesia, followed by the rapid recovery of the female. Extensive details about the mechanical setup, US visualization, and injection procedure are provided in Online Methods.

We successfully applied the FIVI procedure over a broad range of gestational ages (**Fig. 1c**), during which the brain undergoes significant growth and morphological changes. In the marmoset, injections were carried out as early as post-conception day 62 (PC62) (Supplementary Vid. 6 and 7) and as late as PC121. Similarly, in the rat, injections were caried out in fetuses between PC13 (Supplementary Vid. 4 and 5) and PC21, two days before birth. In the earliest injections, the ventricular lumen is more expanded, making it harder to precisely target a specific ventricular compartment. Therefore, we primarily targeted vesicles at the cranial end. For the later injections, the ultrasound-guided procedure allowed more selective targeting of the lateral or 3^rd^ ventricles. Importantly, the capacity to inject at different developmental stages allows for the targeted introduction of viral vectors into specific cell populations as they emerge in the developing brain.

### Widespread rAAV-mediated transgene expression following FIVI delivery

In this section, we describe the widespread viral infection resulting from the FIVI method, revealed through the expression of fluorescent reporter genes (e.g., GFP, tdTomato). Unless stated otherwise, the experiments in this study used the rAAV9 serotype, which has proven effective for injections into developing animals, including the primate fetal brain^23,24^ with reporter expression driven by the high-efficiency CAG promoter (derived from cytomegalovirus, chicken beta-actin, and rabbit beta-globin genes).

Following injections in rats, it was often possible to determine postnatally which of the pups in the litter had received fetal injections by observing reporter fluorescence using a spectrally appropriate flashlight and matching barrier filter glasses (**Fig. 1d**). *In vivo* fluorescence, evident in newborn rats across all gestational injection ages, was visible throughout the head and neck, sometimes extending caudally, and included extracranial labeling of muscle tissue and neurons of the peripheral nervous system. In the marmoset, postnatal *in vivo* evaluation was sometimes possible, though more challenging owing to the pigmentation of the skin, the presence of hair, and the more advanced developmental state at the time of birth relative to rats. When observed, fluorescence in newborn marmosets was most prominent in hairless regions, particularly in the face, around the eyes or in the ears, and inside of the mouth (**Fig. 1d**).

We used histological methods to evaluate the efficiency and cell-type specificity of reporter gene expression in the brain. In general, rAAV9 led to widespread labeling across the cerebral hemispheres. **Fig. 2** shows histological sections of a marmoset injected at PC87 (tissue collected at postnatal day 3) with rAAV9-CAG-GFP, with robust labeling of neurons in lower layers of the cerebral cortex. In this example, we observed dense transduction, with labeling of 30 to 50% of neurons in the labeled layers. Transduced neurons were confined to the lower layers, owing to the relatively early gestational age of injection, discussed in more detail below. In general, a prominent feature of FIVI transduction with rAAV9 is continuous labeling within a laminar band across the cortical sheet (see Extended Data Fig. 1a for additional examples).

**Fig. 2:**
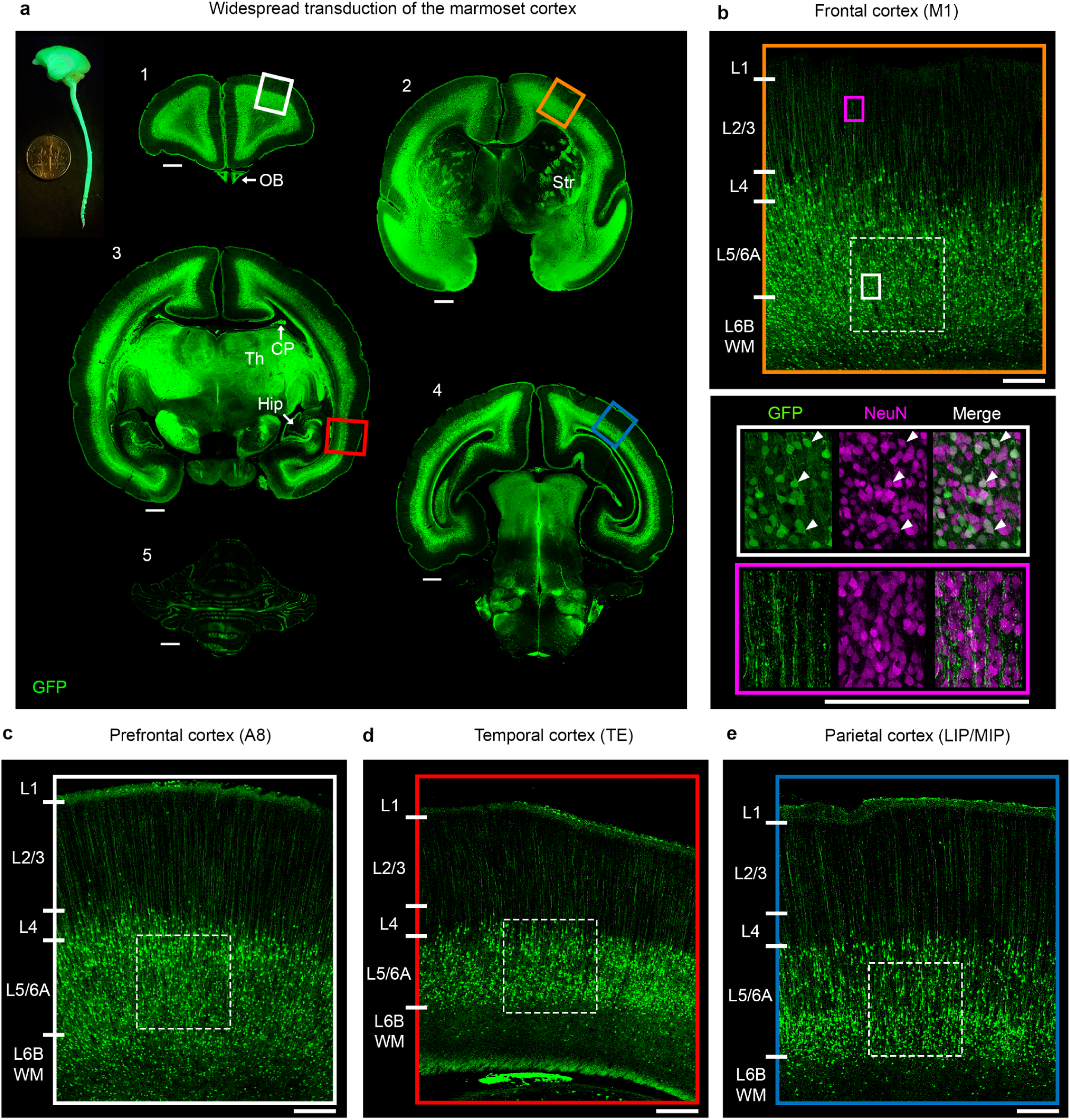
Extensive cortical labeling following FIVI. **a**, Representative coronal sections in a 3-day old marmoset show the widespread transduction pattern following a PC87 injection of AAV9-CAG-GFP. The side view of the extracted central nervous system highlights broad GFP labeling. Across all regions, the most abundant labeling is in lower layers. **b** to **e**, detailed view of the insets shown in **a** highlighting the labeling pattern in the prefrontal, frontal, temporal and parietal regions. Most labeled cells in the cortex are neurons, based on morphology and co-expression of NeuN (panel b, white arrowheads in white inset). Labeling in more superficial layers is virtually restricted to apical dendrites from lower layer neurons (panel b, purple inset). The density of transduced neurons varied between 50 and 30% (dashed squares in panels **b** to **e**) **b**, Frontal cortex, 50% (253/509 GFP+/NeuN+). **c**, Prefrontal cortex, 37% (219/586, GFP+/NeuN+). **d**, Temporal cortex, 30% (275/922 GFP+/NeuN+). **e**. Parietal cortex, 32% (185/586 GFP+/NeuN+). Cell labeling was observed in additional structures, such as the olfactory bulbs (OB, panel a-1), striatum (Str, panel a-2), hippocampus and choroid plexus (Hipp and CP, panel a-3), and cerebellum (panel a-4). Strong fiber labeling was observed in the thalamus (Th, panel a-3) and white matter paths. Scale bars, a: 1 mm; b to e: 250 µm. FIVI injection volume: 60µL.

In rats, the pattern of labeling was comparable to the marmoset. For example, we found similar infragranular labeling (albeit with some additional supragranular cortical neurons) when we injected the same serotype and construct at PC19 (Extended Data Fig. 2a). To evaluate the reliability and reproducibility of this method, we performed the same injection procedure in rat fetuses in 24 sessions, using the rAAV9-CAG construct with injections carried out at PC19 and euthanasia at PC44 (∼21 postnatal days). While some experimental variables, such as the angle of approach to the ventricular system, necessarily varied with each fetus, we fixed other variables. For example, in the PC19 rat, we always injected 10µL of the virus and aimed at the lateral ventricle or midline third ventricle. Although the specific patterns of labeling are subject to the idiosyncrasies of individual injections—for instance, some cases exhibited bilateral symmetrical transduction, while others showed more pronounced expression in one hemisphere—this method reliably led to widespread cortical transfection in multiple fetuses across sessions (Extended Data Fig. 2b and c).

In rats, the expression of reporter genes was first observed within a few days of the procedure (the shortest interval tested was 2 days) and persisted over at least several months (Extended Data Fig. 2d), suggesting this method can be applied to longitudinal neuroscience experiments lasting into adulthood. In marmosets, transgene expression was observed in fetal tissue nine days after the injection (Extended Data Fig. 3). While the maximum duration of transgene expression in marmosets remains unknown, strong expression was observed in adult marmosets 2 years and 4 months post-injection, with no overt neurological or behavioral deficits linked to long-term expression observed in transduced animals.

### Strategies to refine targeting of cell subpopulations

Given the restricted pattern of labeling described above, we investigated factors that might enable targeting different cell subpopulations. One important factor was the gestational timing of the rAAV injection. We compared the FIVI of rAAV9 at four gestational ages in the marmoset ranging from PC85 to PC114 (**Fig. 3a**). We found that the laminar distribution of labeled cells closely followed the injection timing. Earlier injections labeled deeper layers and later injections labeling more superficial layers, consistent with the known inside-out pattern of cortical neuron development (**Fig. 3b**). Thus, adjusting the timing of the FIVI procedure provides a systematic way to target cell populations in the cortex.

**Fig. 3:**
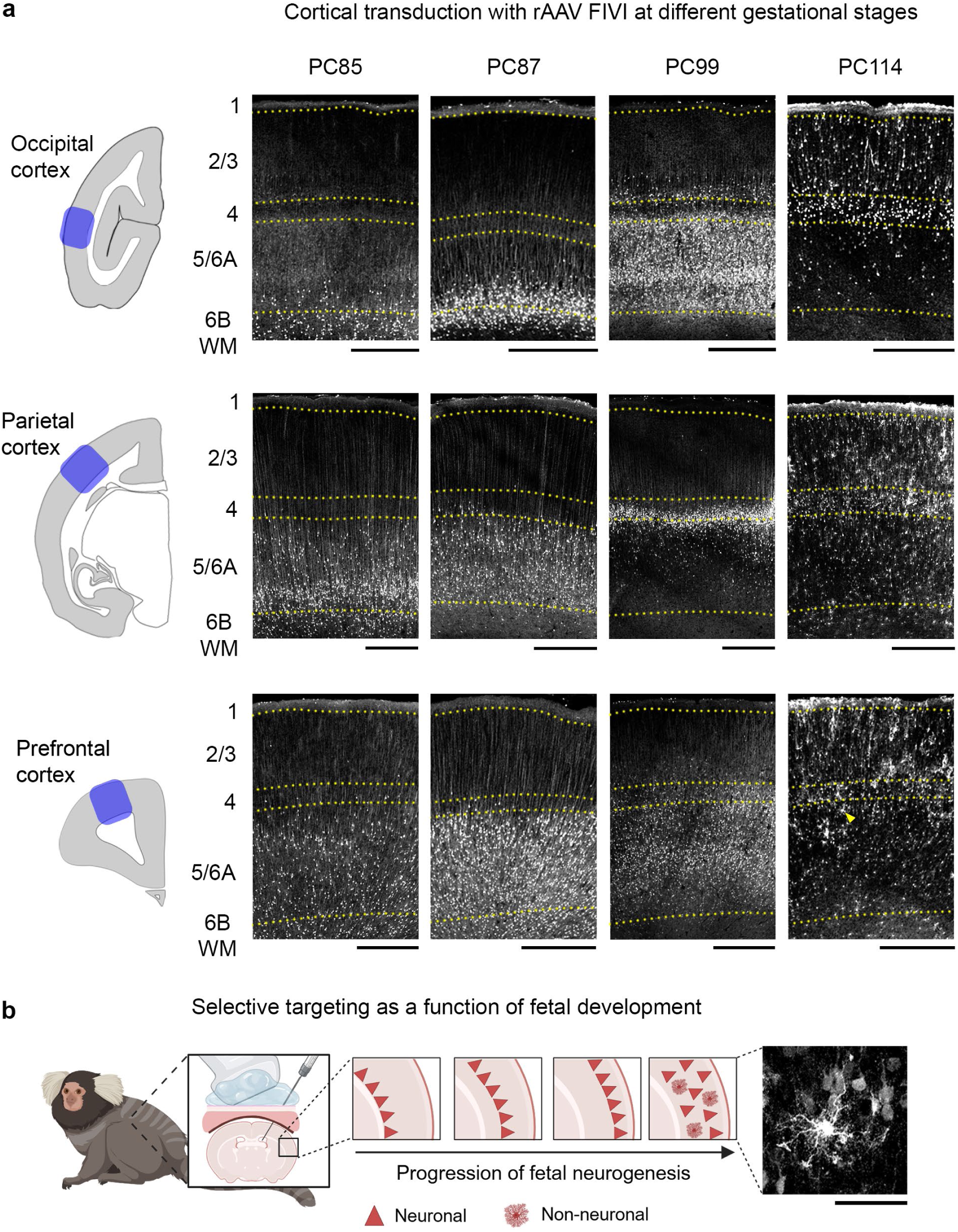
Effects of varying the timing of postconceptional ICV injections. **a**, shows the exemplar perinatal pattern of transduction in the occipital, parietal and prefrontal cortex following rAAV9 FIVI. While there are regional differences, a general trend emerges: earlier injections predominantly label deeper layers, whereas later injections target more superficial layers. This trend is most pronounced in the occipital cortex, where labeling shifts from white matter/layer 6B at PC85 and 87 to layer 5/6B at PC99, and to layers 4 and 2/3 at PC114. For injections administered later in gestation, the labeling differs for parietal and prefrontal cortices. Transduced cells are distributed across all layers and intermingled with non-neuronal cells (arrowhead, non-neuronal cell in panel b histological inset). Tissue collection: PC85/PC98 postnatal day (P) 2; PC87 P3; PC114 P0. **b**, transduction patterns with rAAV9 injections follows the order of cortical histogenesis. Figure panel partially created in BioRender. Scale bars, a: 500 µm; b: 50 µm. FIVI injection volume: 60µL.

The widespread, timing-dependent infection of neurons is influenced by another known property of cortical development, namely that laminar development is asynchronous across the cortical mantle^25^. This can be seen in the different laminar patterns observed in occipital, parietal, and frontal regions (**Fig. 3a**, see Extended Data Fig 1b and 1c for sequential FIVI injections in the same animal). For example, the PC99 injection led to labeling of infragranular neurons in the occipital cortex, but primarily layer 4 neurons in the parietal cortex, with minimal infragranular labeling. Further, the PC114 injection exclusively labeled neurons in the occipital cortex but labeled both neuronal and non-neuronal cells in parietal and frontal areas, suggesting an earlier shift from neurogenesis to gliogenesis in the latter. The interareal differences likely stem from the known rostro-caudal gradient in neurogenesis timing across the cortex^26–29^.

Together, these observations indicate that varying the injection timing within a 30-day interval provides a means to achieve selective laminar transduction across the marmoset cerebral cortex. Furthermore, they highlight a potentially surprising principle of fetal rAAV9 infection in primates: neural labeling appears to be confined to cortical cells born within a window of time relative to the injection procedure. While the absence of labeling superficial to the densely labelled band can be explained by upper-layer cells being born or reaching a critical maturational state permissive for rAAV transduction after the injection, the absence of deeper labeling is more puzzling. This absence suggests that some cells born *prior* to the procedure were no longer eligible for transduction. While the reason is unknown, it may stem from the physical position occupied by those radially migrating neurons at the time of injection. Alternatively, it may be due to a change in the cell surface molecules associated with normal maturation, such as the specific membrane glycoproteins known to act as co-receptors for rAAV infection^30^.

Thus, one possibility is that the selective transduction of neurons based on maturation stage derives from viral tropism, where only neurons at a given maturational stage are transduced by a given rAAV serotype expressing particular capsid proteins. To investigate this possibility, we compared the patterns of infection with two different serotypes, rAAV2 and rAAV9. Pilot studies had revealed these two serotypes to exhibit different laminar expression patterns in rats when injected at the same gestational age (Extended Data Fig. 2e). When rAAV2 and rAAV9 were injected into marmosets at comparable gestational ages using the FIVI method, we observed a pronounced serotype-specific laminar segregation across the cortex (**Fig. 4**). Namely, rAAV9 labeled the earlier born, deep layer neurons, whereas rAAV2 labeled the later born, superficial layer neurons. This observed serotype-driven laminar transduction is consistent with an interaction between the intrinsic tropism of the virus and the developmental state of the cells.

**Fig. 4:**
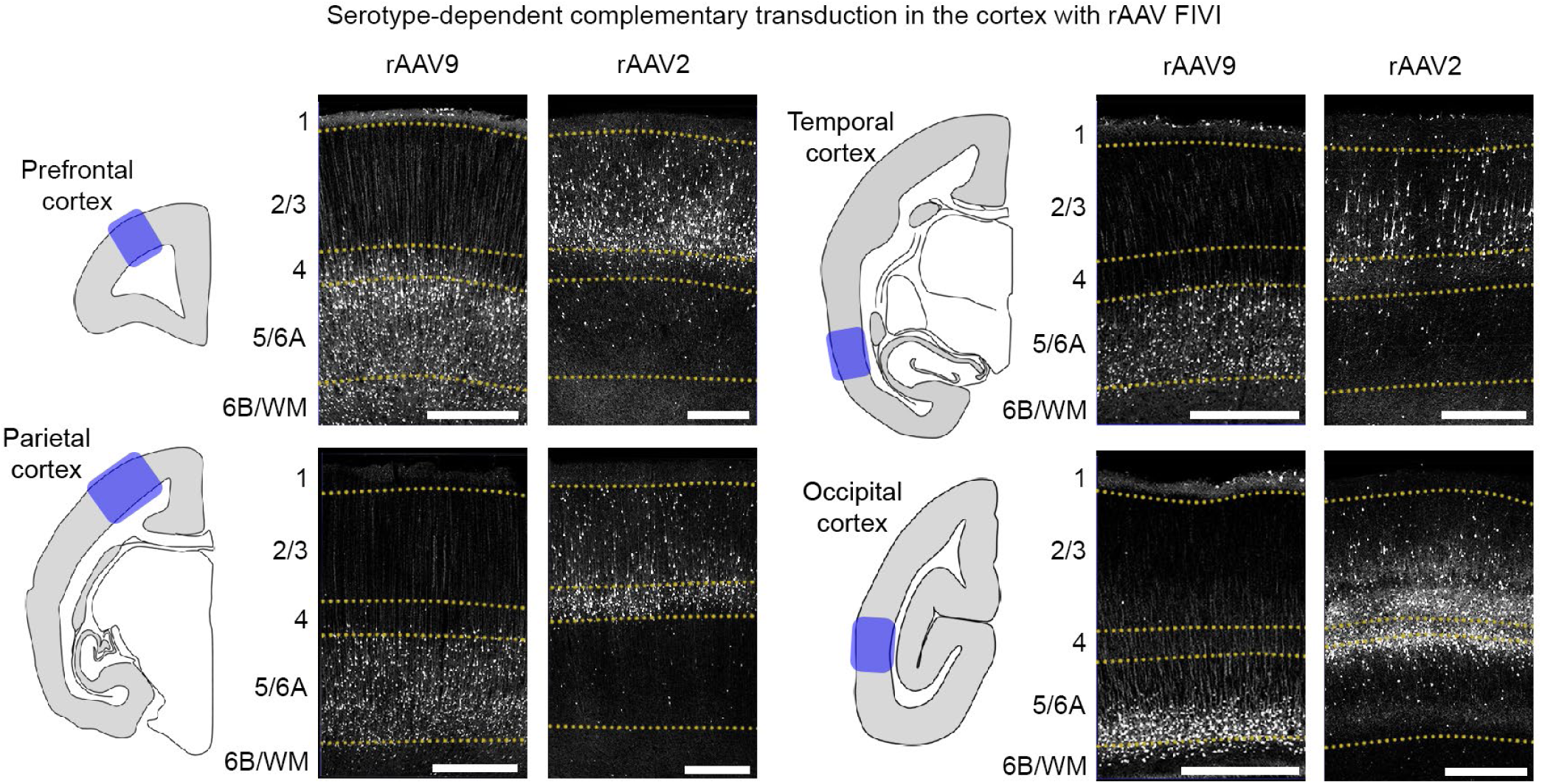
Viral tropism-mediated selective transduction of the fetal brain. Representative perinatal transduction patterns in the prefrontal, temporal, parietal, and occipital cortices of the marmoset following rAAV2 and rAAV9 FIVI at similar gestation times. As in rats (Extended Data Fig. 2e), the serotypes exhibit complementary transduction patterns, with rAAV2 labeling neurons in more superficial layers compared to rAAV9. Notably, as shown in Fig. 3, the transduction patterns vary across regions. Tissue collection: rAAV2 PC86 at P2; rAAV9 PC87 at P3. Scale bars, 500 µm. FIVI injection volumes: 60µL.

Finally, for neurodevelopmental studies, the FIVI methodology provides a powerful tool to observe and manipulate cells as their identity is established and cell diversity emerges^31^. For example, by employing rAAV intersectional strategies in marmosets, FIVI can be used to tag cells based on the transient prenatal expression of a specific gene. We applied this rationale to the *Nestin* gene, which encodes an intermediate filament protein expressed in neural stem cells as well as subsets of non-neural stem cells^32^. It is also known that Nestin expression progressively declines shortly after cells differentiate^33^. Using FIVI procedures at different gestational ages in two marmosets, we co-injected two viral constructs to tag neurons expressing Nestin at or after the injection procedure. The first vector (AAV9-NestinTK-EGFP-iCre) expressed Cre recombinase under control of a Nestin-specific enhancer in undifferentiated cells^34^. The second vector (AAV8-nEF-Con/Foff 2.0-ChRmine-oScarlet), injected at the same time, led to the Cre-dependent expression of the fluorescent reporter oScarlet in Cre-expressing cells. With this combination, it was possible to identify cells that exhibited Nestin expression after either PC105 (Extended Data Fig. 4b and c) or PC86 (Extended Data Fig. 4d). This proof-of-concept intersectional experiment demonstrates the potential power of the FIVI method to identify and study developmentally defined neural subpopulations across the primate brain.

### Prenatal gene editing using the CRISPR/Cas9 toolbox

The precise editing of a cell’s genome is an important advance in biology, with enormous prospects for human medicine^35^. For somatic tissue, including the embryonic brain, gene editing can be achieved using the CRISPR/Cas9 toolbox delivered by rAAV. To this end, we modified constructs previously validated in mice^36^ to insert the reporter EGFP into the rat *Actin* gene to create an EGFP-Actin fusion protein through CRISPR/Cas9-mediated homology-directed repair. This gene editing in rats was achieved by co-injecting a vector expressing SpCas9 (AAV9-EFS-SpCas9) along with a vector expressing a guide RNA targeting the start codon of rat *Actin* as well as carrying a repair template (the coding region of GFP flanked by homologous arms specific for sequences flanking the CRISPR induced double strand break) (AAV9-rActin-EGFP-donor) (**Fig. 5a**). Applying this two-virus approach resulted in EGFP expression among a sparse subset of neurons throughout the brain following FIVI at PC19 (**Fig. 5b**). These neurons exhibited EGFP signals localized in known Actin-rich regions, such as the dendritic spines (panels D and E). EGFP expressing cells were also observed in the choroid plexus (panel F), a tissue susceptible to rAAV infection and a potential source for widespread delivery of biotherapies to the brain, as well as in other regions amenable to rAAV9 serotype transduction at PC19.

**Fig. 5:**
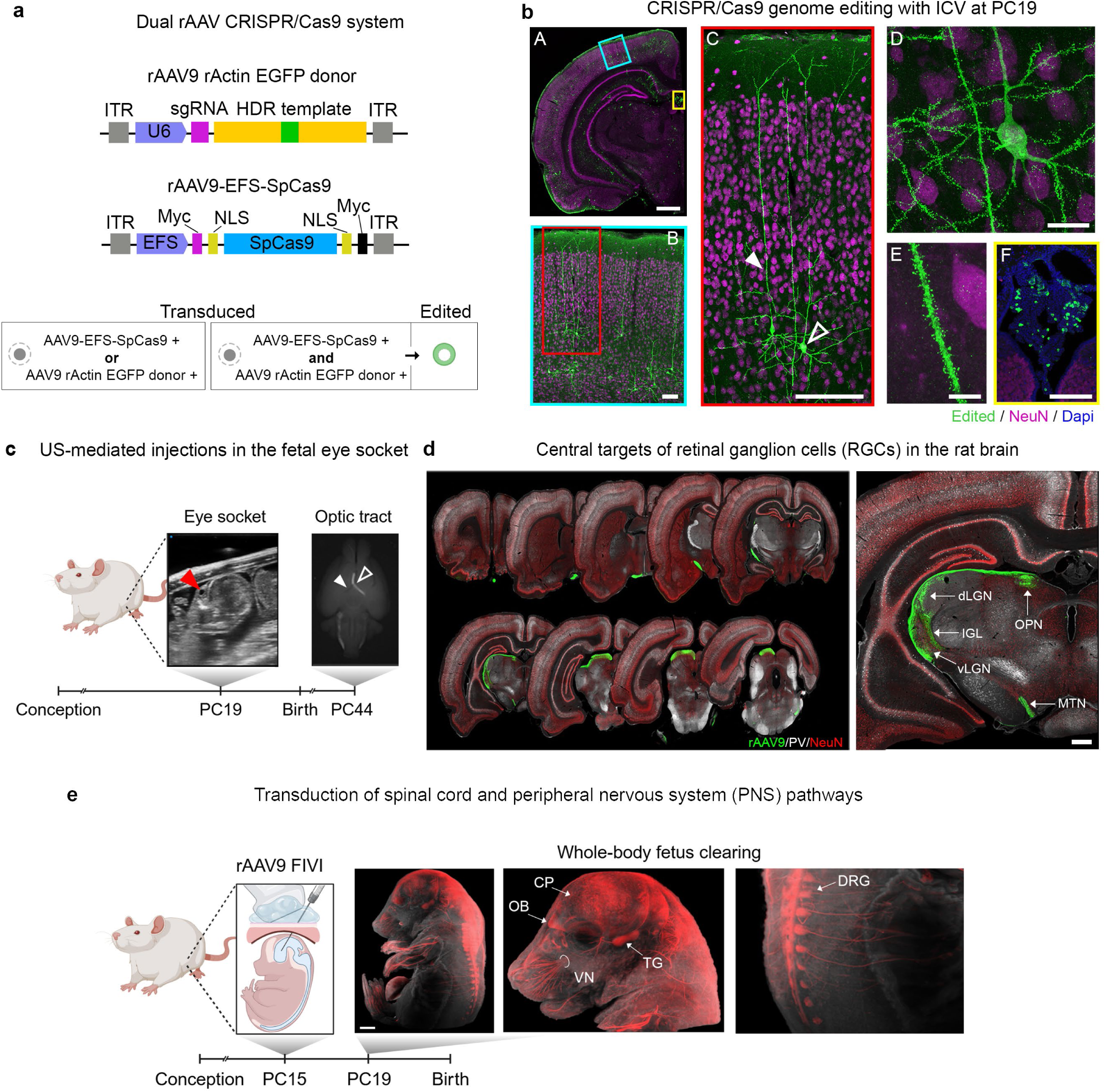
Proof of concept gene-editing and peripheral-labeling experiments using FIVI. **a**, Gene editing with CRISPR/Cas9. CRISPR/Cas9-mediated genome editing in the rat through co-transduction with rAAV9-EFS-SpCas9 (5µL) and rAAV9-rActin-EGFP-donor (5µL). **b**, FIVI at PC19 with dual rAAV CRISPR/Cas9 system led to sparse gene editing of neurons in the cerebral cortex at PC44 (B and C). The EGFP reporter fused to actin fills the neurons and accumulates in dendritic spines such as those highlighted in panels D and E. F highlights edited cells in the choroid plexus. Scale bars, A: 1 mm, B: 500 µm, C: to E 10 µm. Diagram adapted from Nishiyama and colleagues^35^. **c**, transduced retinal ganglion cells (RGCs) labeled with a PC19 rAAV9-CAG-GFP (2µL) injection into the right eye socket (red arrow). RGC axons decussate in the chiasma (open arrowhead) and project to targets in the contralateral hemisphere. Note the lack of label in the contralateral optic tract (closed arrowhead). **d**, Sequential coronal sections (rostral to caudal) at PC44 showing the pathway and the central targets of RGCs projections, such as the dorsal lateral geniculate nucleus (dLGN), the ventral LGN (vLGN), the intergeniculate leaflet (IGL), the medial terminal nucleus (MTN), and the olivary pretectal nucleus (OPN). Scale bar, 500 µm. **e**, FIVI injections lead to both central and peripheral nervous system labeling. Optical clearing of a PC19 transduced fetus, injected at PC15 with rAAV9-CAG-tdTomato (5µL), enables 3D visualization of reporter gene expression in the central (e.g., olfactory bulbs (OB), cortical plate (CP), spinal cord) and peripheral nervous system (e.g., dorsal root ganglion (DRG), trigeminal ganglion (TG), vibrissal nerves (VN)). This approach facilitates investigation of the spatial and temporal development of cell type specific projection pathways. Figure partially created in BioRender.

### Prenatal transduction of the peripheral nervous system and sensory pathways

Beyond labeling cells in the central nervous system, the FIVI approach can be used to transduce peripheral neurons and sensory pathways, as we illustrate with two examples, both of which use rAAV9 and a fluorescent reporter under the control of CAG. The first example, shown in **Fig. 5c** and **5d**, entails the selective labeling of retinal ganglion cells (RGCs) in rats at PC44 following ultrasound-guided targeting of the eye socket at PC19 with rAAV9-CAG-GFP. A single ocular injection led to prominent transduction of RGCs projections exiting the eye, highlighting the many visual ganglion cell projection sites in the brain. Combined with more sophisticated viral approaches, this paradigm may be used to study the early development and childhood refinement of retinal pathways, including in nonhuman primates. The second example, shown in **Fig. 5e**, illustrate the rAAV9-CAG-tdTomato labeling in the peripheral nervous system following a PC15 injection. In this case, the fetus was harvested four days later, at PC19, and histologically cleared. In the cleared fetus, tdTomato labeling was prominent in the dorsal root ganglia, trigeminal ganglia, and peripheral nerves. Across FIVI experiments transduction of spinal cord and sensory neurons was a common observation, in agreement with previous studies^37^.

## Discussion

This study describes a minimally invasive ultrasound-guided fetal intracerebroventricular delivery method offering new experimental and translational opportunities for widespread viral gene transfer, particularly in primates. We used the rat model to establish and refine the FIVI setup and procedure, as well as to compare transgene expression patterns with marmosets. In this initial study, we focused primarily on rAAV9, as this serotype has been shown to lead to efficient transgene expression following direct injection into the primate brain^20,38^, including during early development^24,39,40^.

Extensive reporter expression following FIVI was prominent in the cerebral cortex. The cortex-wide transduction paves the way for a range of neuroscience experiments facilitated by broader expression of opsins^41^, calcium indicators^42,43^, or other genetically encoded neuroscience tools. Investigations utilizing these methods, firmly established in mice, can now be translated to primates, particularly the marmoset with its lissencephalic cortex. Since transgene expression starts during fetal development, this emerging combination of methods is well suited for longitudinal studies offering novel prospects for mapping whole-brain circuits and tracking physiological or behavioral changes across the lifespan. Outside the cerebral cortex, transduction of other regions depended on the specific combination of rAAV serotype and on the injection timing relative to gestation. Refining and optimizing transduction targeting with the FIVI, including in non-cortical structures, remains a key focus for future studies.

It is useful to compare rAAV delivery with FIVI with other *in vivo* approaches. Intraparenchymal injections have become a standard method in primate neuroscience^44–46^. Typically applied in adults, local injections allow for spatial restriction of gene transfer to regions of interest, due to the relatively short diffusion distance. While this facilitates precise targeting with high levels of transgene expression, it excludes experimental opportunities that come from widespread gene transfer across the entire brain. An emerging approach in adults for broader expression is the intravenous infusion of rAAV9. This strategy was initially based on the capacity of this recombinant serotype^47–49^ and later enhanced by its capsid-engineered variants^50–52^ to efficiently cross the blood-brain barrier and infect neural tissue. Successfully applied in nonhuman primates^24,53–55^, intravenous delivery methods are conceptually straightforward. With improvements in vector design to decrease off-target transduction and immunological response, these approaches are becoming more commonly used. While holding promise for postnatal transduction, these methods have limited prenatal application, particularly for fetal transduction following infusions in the pregnant dam. To our knowledge, this approach does not lead to fetal transduction, likely due to the inefficiency of the capsids in crossing the placenta.

For prenatal studies, electroporation methods have been the most widely used tools for in vivo gene transfer for neurodevelopmental studies. It involves the spatially restricted introduction of DNA into neural precursor cells through a surgical procedure exposing the fetus^56,57^. While firmly established in the mouse, electroporation is highly invasive, less effective in later stages of development, and achieves lower rates of expression compared to viral vectors^58^. For primate experiments, there are high risks for both fetus and dam associated with a major surgical procedure involving the exposure and manipulation of fetuses. Though the outcomes of these procedures can be improved^59–61^, risk mitigation requires specialized training and large numbers of animals, thus severely restricting the number of research groups that could take such an approach. The FIVI method offer a more tractable solution to these challenges, as well as a number of unique advantages. Importantly, by enabling safe and flexible delivery of research and therapeutic products using viral tools or other emerging therapeutic vehicles^62,63^, it keeps pace with rapid advances in delivery technologies, enabling rapid evaluation and selection of effective tools for both fundamental and translational research. FIVI will facilitate the investigation of intricate mechanisms underlying neurodevelopment such as cell history, fate, and specification, while also enabling new opportunities for study interventions during critical windows of prenatal development. This has the potential for long-lasting and efficient gene modulation, informing on long-term effects of prenatal transduction, valuable information for disorders where *in utero* intervention can lead to improved outcomes^64^. For restorative gene therapy, direct editing of the chromosomal DNA using methods such as CRISPR may prove most effective^35,65^. The transduction of not only neurons, but also important cellular targets for gene therapy such as cells in the choroid plexus^66–68^, suggests that the FIVI procedure can be a viable approach for therapeutic applications.

It is also important to acknowledge limitations of the current method and potential areas for future improvements. One key challenge in the application of viral vectors is the control over the specific tissues and cells being target and transduced. High cellular tropism specificity, with targeting of a narrow range of cell types, can be either desirable or limiting, depending on the intended transduction goal^69^. The examples shown in the present study demonstrate both sides of this issue. For the broad entry and transduction of rAAV across multiple cell types, followed by the restriction of transgene expression based on promoter or enhancer sequences, the tropisms observed with FIVI, such as the incomplete coverage of the cortical mantle, limit the extent to which all cell populations can be uniformly transduced. On the other hand, this method provides an additional strategy for cell selection based on combinations between gestation timing and serotype used, as observed in the complementary laminar patterns of neurons labeled at different times (**Fig. 3**) or by two different serotypes (**Fig. 4**). Significant gaps remain in our understanding of rAAV transduction in primates, particularly regarding the parameters that influence transduction efficiency. For instance, intracortical injections of nine different serotypes (AAV1, 2, 5, 6, 7, 8, 9, rh10, and DJ) at moderate titers predominantly resulted in transgene expression in neurons when using the ubiquitous CMV promoter^70^. However, a fourfold increase in titer led to a shift from neuronal to glial transduction^71^. This gap in knowledge is especially pronounced in fetal brains. It is clear that the rAAV transduction patterns observed in this study differ significantly from those observed in adults. A possible explanation for this difference may be differences in the composition of rAAVs binding sites in target cells during their maturation. Glycans, such as proteoglycans^72^, serve as AAV receptors and play a crucial role in various mechanisms underlying nervous system development and maturation. Their expression is tightly regulated both spatially and temporally during development^73^. Future studies on the spatial and temporal dynamic changes in the developing glycome^74^ may enhance our understanding of the host cell-rAAV binding, which may be at the origin of differential transduction patterns. This knowledge will be valuable for designing experiments to optimize rAAV transduction based on the cell’s developmental state, or for transducing cells at complementary stages (e.g., by combining serotypes), thereby expanding the coverage of gene transfer.

One potential concern with the fetal injections is the risk of inducing abortions or otherwise affecting the pregnancy or development of the fetus. In our case, while abortions did occur in the marmoset, they were rare and did not immediately follow the procedure. As spontaneous abortion is common in marmosets, the reproductive losses we observed in injected females were on par with those described in the literature^75^ and were concomitant with miscarriages in non-injected females in our colony. More likely explanations, such as environmental stressors (e.g., abrupt reconfigurations in the marmoset’s environment) ^76^ and litter size^75^, cannot be excluded.

Another potential concern with fetal injections is the possibility of morphological changes in brain development following FIVI. These could result from the interaction of multiple factors, including the FIVI methodology (e.g., anesthesia duration, intracranial targeting), parameters of rAAV delivery (e.g., needle gauge, injection pressure, dose and volume), or characteristics specific to the injected subject (e.g., gestation timing, immunological response). We occasionally observed morphological changes (e.g., dilated cerebral ventricles and modifications in the typical sulci and gyri morphology) in the injected brains of rats and marmosets. For instance, enlarged ventricles were observed in 2 of the 11 brains analyzed histologically, both in brains that received sequential injections of rAAV (see section of Fetus 3 in Extended Data Fig. 1a for enlarged ventricle example). These brains also presented variations in the typical sulci and gyri morphology (e.g. increase in the deepness of the posterior occipital sulcus (pos) and the presence of a sulcus in the inferior posterior occipital cortex in Fetus 2 in Extended Data Fig. 1a). Whether morphological variations result from the known variability in individual marmoset brain morphologies^77–79^ or are accentuated by biomechanical stress induced by the procedure or the genetic manipulation itself remains unclear and requires further investigation. Exploring the use of smaller injection volumes scaled proportionally to each fetus may help mitigate these changes by reducing CSF pressure and preserving CSF chemical balance. Investigating the use of lower titers, promoters with lower transcription rates, or inducible viral systems may also be considered if adverse effects of strong and long-term exogenous gene expression on cellular function are suspected. Diversifying delivery methods, such as ultrasound-guided retroorbital injections or injections into the fetal circulation, could provide complementary routes for fetal transduction. Optimizing rAAV vectors, of recombinant or engineered origins, with increased transduction efficiency in progenitors^80^, may broaden the extent of transduction, and help circumvent the need for repeated rAAVs administrations to reach sufficient expression levels (e.g., for therapeutic effect). Further refinement of the methodology could improve targeting consistency, shorten the procedure, decrease anesthesia time, and minimize complications that may arise with longer procedures.

Finally, additional studies are necessary to assess whole-body prenatal biodistribution of rAAVs and the susceptibility of the immature primate brain to these vectors. Recombinant AAVs such as those used in this study have been shown to have variable efficiency in transducing dividing cells^81^. Assessing their efficiency in transducing dividing cells *in vivo* during primate development and their impact on the proliferative capacity and survival, will help select serotypes that may be safer for prenatal transduction at different gestational stages. Further studies investigating the immunological tolerance or ignorance of the fetal brain toward rAAV or transgene products will help refine protocols for fetal transduction (e.g., increasing the interval between sequential injections if these are required). Longitudinal monitoring of neurological function and behavior from birth will provide valuable insights into optimizing parameters (e.g., titer) for safe rAAV administration.

In summary, we presented a novel method for prenatal *in vivo* delivery of genetic technologies in nonhuman primates, emphasizing critical factors influencing rAAV vector transduction, such as gestational timing and rAAV serotype. The versatility of FIVI, which allows for routine testing and troubleshooting, holds significant potential for establishing best practices and successful protocols for *in utero* gene transfer. Its broad applicability across species, and particularly in primates, will further support the development of additional quasi-transgenic animal models for experimental and preclinical research.

## Acknowledgments

This work was supported by funding from the Intramural Research Program of the National Institute of Mental Health (ZIAMH002898) to D.A.L and National Institute of Child Health and Human Development (P50HD103536) University of Rochester Intellectual and Developmental Disabilities Research Center (UR-IDDRC) to K.H.W. ARRG was funded by the National Institute of Health Visiting Fellow Intramural Research Training Award. NH and NW were funded by the National Institute of Health postbaccalaureate Intramural Research Training Award. We thank George Dold and members of the Section on Instrumentation of the Intramural Research Program of National Institute of Mental Health for their support in designing and fabricate custom-made parts used in this project. We are grateful for support from the Systems Neuroscience Imaging Resource of the Intramural Research Program of the National Institute of Mental Health for their allocation of imaging resources used in parts of this research. We thank the Genetic Engineering and Viral Vector Core from the Intramural Research Program of the National Institute of Drug Abuse for the development and production of genetic tools used in this study. We also thank the Veterinary Medicine and Resources Branch from the Intramural Research Program of the National Institute of Mental Health for their support with veterinary health care, animal husbandry, and technical assistance. Finally, we are grateful to Lenegereshe Baweke and Sean Kearney for technical assistance.

## Online Methods

All procedures were approved by the Animal Care and Use Committee (ACUC) of the Intramural Research Program of the National Institute of Mental Health (NIMH) and performed at the U.S. National Institutes of Health in accordance with institutional guidelines.

### Animals

Time pregnant Sprague Dawley® rats (SD strain) were acquired from Charles River Inc. They were singly housed upon arrival with gestational ages varying between postconceptional (PC) day 3 and PC10 and acclimated to the vivarium for at least 72h before the procedure. Postconceptional day 1 corresponds to the day when an ejaculatory plug was observed. All rats were housed in a temperature and humidity-controlled environment, under 12h/12h reversed light/dark cycle with ad libitum access to food and water.

Fetal injections were done in sixteen adult common marmoset (Callithrix jacchus) females, with ages between 2 and 9 years old. All animals were paired and housed in family groups, with ad libitum access to food and water, and in a temperature and humidity-controlled environment with 12h/12h light/dark cycle. Marmoset dams were reinjected in successive pregnancies without an increased risk of miscarriage or fetal death. The histological data in this report is from the offspring of six.

### Marmoset ultrasound pregnancy evaluation and estimation of gestational day

Marmoset pregnancies were detected by transabdominal ultrasonography and monitored at least once monthly using the FUJIFILM VisualSonics Vevo MD UHF22 and UHF48 linear transducers. The females were hand restrained to minimize movement, and given a positive reward (dried fruits or pediasure) before, during, and after each scan. The ventro-dorsal and transverse diameter of the uterus and uterine lumen, along with the biparietal diameter of fetus skull were measured. The PC dates for the procedures were estimated using the published values from Oerke and colleagues^1^. Subsequently, these dates were re-estimated after birth by subtracting 143 days from birth, accounting for the average pregnancy length in marmosets, between 140 and 145 days ^1^. The difference between the estimated gestational day for each female before the procedure and the day after delivery varied between 0 and 9 days. The median difference between PC days estimated by fetal ultrasound and post-delivery estimates is 0, while the median absolute difference is 3 days, indicating good agreement between the two approaches. The dates used in this report are the dates re-estimated following delivery, except for the PC114 case reported in Fig. 3. This neonate was delivered via c-section at the estimated date of birth under veterinary guidance due to a history of dystocia in the previous pregnancies.

### Viral constructs

All the constructs used in this study were from recombinant Adeno-Associated Viruses (rAAV) serotypes. The rAAV plasmids or viral preparations were acquired from Addgene or prepared by the National Institute on Drug Abuse Genetic Engineering and Viral Vector Core Facility (RRID:SCR_022969). Details on the viral vectors, including their sources and the investigators who gifted the items, can be consulted in Supplementary Table 1.

### Plasmid construction and vector packaging

Details on the plasmids, including their sources and the investigators who gifted the items, can be consulted in Supplementary Table 2. All constructions were performed with ligation-independent cloning (In-Fusion, Takara), and transformed into NEB Stable cells. Isolated colonies were miniprepped and verified by fragment analysis and sequencing prior to viral packaging.

The plasmid pOTTC2234 (RRID:Addgene_228443) was constructed by amplifying the sequence corresponding to the human nestin 2nd intron enhancer with TK mini promoter and EGFP (Nestin-TKmini promoter-EGFP) from Addgene 38777 (RRID:Addgene_38777) and the sequence corresponding to iCre from the plasmid pOTTC1031 (pAAV CMV-IE eGFP-2A-iCre), followed by their insertion into a plasmid already containing a WPRE regulatory element, a TK65pA poly-adenylation signal, and two inverted terminal repeats (ITRs) for AAV (pOTTC2210) ^2^. The open reading frames of EGFP and iCre are separated by a 2xglycine linker.

The plasmid pOTTC2360 (RRID:Addgene_228444) was constructed by amplifying the nucleotide sequences upstream (-971 to -1 bp) and downstream (4 to 1014 bp) with respect to the rat actin start codon and using them to replace the homology arms corresponding to mouse actin in pAAV-HDR-mEGFP-Actin (RRID:Addgene_119870) ^3^. The guide RNA expression cassette was also retargeted from mouse actin to rat actin using the following seed sequence (TGTGCCTTGATAGTTCGCCA).

AAV viral vectors were packaged using calcium phosphate precipitation to transfect HEK293 cells with a mix of the pAAV genomic cargo plasmid and two trans plasmids (pHelper and pAAV 2/9 (p0008) as previously described^4^. Transfected cell pellets were thawed, then freeze-thawed two times, vortexing after each thaw. MgCl2 (2 mM final; Sigma-Aldrich, St. Louis, MO) and SAN HQ (400 U/mL final, Arcticzymes, 70920-150) were added to the cell solution and incubated at 37°C with continuous shaking for 1 hour. The solution was centrifuged for 20 min at 2450 × g at 4°C. The supernatant was transferred to 75 ml of phosphate-buffered saline (PBS) with 2 mM MgCl2 and sequentially filtered through 5-, 0.45-, and 0.22-μm filters. This crude solution was then run through a 1 mL POROS GOPURE AAV9 Column (Life Technologies/Invitrogen, A36650) using an AKTA Pure25 FPLC (Cytiva) at a rate of 2 mL/min and eluted using a solution of 200mM Sodium citrate (pH2) + 400mM NaCl (Sigma-Aldrich) at a rate of 1mL/min. The fractions containing the peak of the UV absorbance (A254 and A280) were collected and dialyzed using a 10,000 MWCO dialysis cassette (Pierce, now Thermo Fisher Scientific, Rockford, IL) in 1 L of PBS containing 0.5 mM MgCl2 for three exchanges over 25–30 hours. The equilibrated virus was aliquoted, snap-frozen, and stored at −70°C. Purified vectors were titered by droplet digital PCR with a probe-based assay that recognizes the EGFP or SpCas9.

### ICV injection setup

To successfully perform fetal intracerebral ventricular injections (FIVI) without the need for surgery, we designed a system allowing repositioning of the pregnant dam, for better visualization and targeting of the fetuses, while maintaining stability and reducing fetal movements throughout the procedure. This system is fully mechanical, and most of its parts are commercially available. Only a few elements were custom designed by the NIMH Section on Instrumentation, such as the guide tube, transducer holders and the cradle. Drawings for these parts are available upon request. Details of the FIVI setup components are listed in Supplementary Table 3.

Figure 1a illustrates the assembly of the setup, highlighting its key components, as well as the general layout we use in our procedures, in which the different items are set up to maximize the use of smaller spaces. All supplies and equipment are portable, and their positioning depends on the specific procedure and the room in which it is carried out. All procedure rooms are equipped with a benchtop or a surgical table to place the setup and the electrosurgery unit and has enough space to accommodate a portable anesthesia machine, monitoring systems, and an ultrasound machine. The ultrasound machine is positioned near the person performing the injections. A main component of the FIVI setup is the cradle, composed by a 25 cm length hemicylindrical tube of 15 cm width. At one end of the cradle, a Loc-Line® adjustable arm was connected to a stainless-steel extended spring clip for flexible, but stable fixation of the anesthesia mask during the procedure. In both rat and marmoset procedures, for inducing and maintaining anesthesia, we used a RC2 rodent circuit controller (VetEquip), fitted with a nylon-reinforced, 0.25-inch inner diameter, conductive gas supply hose for wall oxygen supply. Since this is a short procedure, oxygen can also be delivered using a high-pressure cylinder. The cradle is supported by a cube geared tripod head (Arca-Swiss C1), clamped to the cradle via a 3D printed dovetail rail fixable to the quick release clamp from the geared head. The length of the rail encompasses the full length of the cradle. This facilitates the initial positioning of the animal beneath the stereotaxic arms and allows for rapid repositioning of the pregnant dam over longer centimeters distances (e.g., quick shift between fetuses in the upper and lower abdominal quadrants). Depending on the layout of the room and dexterity of the person doing the injections, the cradle can be positioned with the head of the animal to the right or to the left. The 62° tilt-only axis of the geared head enables rotation of the cradle and allows fine adjustments to the fetus orientation. The geared head is mounted on two single axis translation stages, mounted on a breadboard. The bottom stage is used for x-axis translation and, and the top stage, rotated 90° relative to the bottom stage, was added for y-axis translation. These plates are equally useful for slower millimeter to centimeter distance adjustments, such as when moving from one rat fetus to another in a neighboring sac within the same uterine horn, or when readjusting the needle trajectory before and after penetration. Two handles were added to the breadboard base for easier transport.

The stereotaxic arms and micromanipulators are carried by an A/P bar attached to a travel vertical translational stage, which allows additional vertical adjustments. The placement of the stereotaxic arms behind the animal gives easy access to the abdomen for the injections. The approximate 8-centimeter space between the geared head and the bar allows the cradle to *float* without touching the A/P bar. This gives sufficient leeway for cradle rotations and y-axis adjustments. We found that using a cradle, rather than a flat surface, was the best design for stabilizing the animal while moving and when rotated, eliminating the need for taping the animal. It also improves temperature maintenance provided by the heating pad throughout the procedure.

The guide needle and ultrasound holders are caried independently by two rotation adapters, fixed to an anterior-posterior (a.p.) slide attachment mounted onto stereotaxic arms. The rotation adapter carrying the ultrasound transducer holder, 3D printed and mounted on a stainless-steel rod, is used to angle the transducer. This provides space for positioning the guide beneath the transducer, allowing it to be visualized on the ultrasound images (Supplementary Vid. 1 and 2). The rotation adapter carrying the guide holder allows slight rotations of the guide. This feature is sometimes useful for correcting the injection trajectory after skin penetration, without the need to modify the angle of the stereotaxic arm. The a.p. slide attachments are used to further adjust the guide relative to the ultrasound transducer. Finer, millimeter or less, a.p. adjustments are achieved using the micromanipulators supporting the stereotaxic arms.

To penetrate the skin and inject the cranium, we used custom-made Hamilton Company needles. For the guide tubes we used 23-gauge or 24-gauge needles. To concentrate the small electrical current delivered by the electrosurgery unit to the tip of the guide, the guides were sheathed with Palladium™ Pebax™ Heat Shrink Tubing using a heat gun. The blunt and beveled ends of the guide tube were uncoated. The guide tube was mounted on a custom-made holder, made of two anodized aluminum plates mounted on a stainless-steel rod. The guide is inserted into a small indentation between the two aluminum parts and secured in place by tightening the lateral screw. The current produced by the electrosurgery unit is transferred to the guide tube using a pair of alligator clamps. One end connects to the blunt end of the guide, and the other to the electrosurgery pencil. Using current greatly facilitates penetration of the skin (Supplementary Vid. 1 vs 2).

To inject the fetuses, we use 31-gauge to 33-gauge small hub removable needles. Depending on injected volume, we used 5, 10, 50 or 100µl Model 701 Hamilton syringes.

### Injection procedure and ultrasound-guided injections

Before the procedure, the needle guide holder, forceps, guides, and needles were cleaned, disinfected, and sterilized with ethylene oxide. Unless bent, needle guides and injection needles were reused across procedures. At the start of the procedure, the ultrasound transducer was disinfected using germicidal disposable wipes (Sani-Cloth® Prime). The room was divided in two spaces, one for preparing the animal and one for doing the injections. FIVI in rats were made between PC13 and 21, while injections in fetal marmosets were done between PC62 and 121. At the start of the procedure, pregnant rats received a dose of meloxicam (1 mg/kg; 1.5 mg/ml (p.o.)) for preemptive analgesia, and pregnant marmosets received both meloxicam (0.1-0.2 mg/kg, 5 mg/ml (p.o., i.m or s.q.)) for preemptive analgesia and diazepam (0.25-1.0 mg/kg, 5 mg/ml (i.m)) for light sedation. Note that for marmosets, the procedure on pregnant dams was performed on after fasting to minimize the risk of complications associated with regurgitation and aspiration under anesthesia. Following premedication, the dam was anesthetized with isoflurane gas (2-3% induction, 1-2% maintenance), and transferred to the cradle in the prep area. Ophthalmic eye ointment was then applied to both eyes to prevent dryness. The abdominal fur was trimmed and then shaved with depilatory cream (Nair cream) to remove hair and enhance ultrasound visualization of the fetuses. A small patch of fur (about 2 x 3 cm) was trimmed on the back for electrode grounding. After removing all hair clippings, the dam was gently placed in the supine position over the electrosurgery grounding pad in the cradle and transferred to the injection setup. Non-sterile ultrasound gel was applied between the trimmed back and the dispersive electrode to improve contact and to safely return the electrosurgical energy delivered through the guide, minimizing damage or burns to the female. The lateral sides of the pad are taped to the heating pad to secure the animal from sliding out of the cradle with larger angles. To prevent possible pressure on the inferior vena cava due to the weight of the uterus, the cradle was angled to elevate the head. Throughout the procedure, the animal’s respiratory rate was manually monitored every 10 minutes. Additionally, a physiological monitoring system (rat: Kent Scientific SomnoSuite®; marmoset: IntelliVue MX500) was used to monitor the animal’s pulse oximetry, and to monitor and regulate body temperature. In marmosets, the system further allowed for monitoring of expired CO_2_, and continuous recording of respiratory and heart rates. In both species, supplemental heating was provided using a benchtop infrared heater and an overhead heating lamp to ensure regulation of the animal’s body temperature around 37°C and prevent hypothermia. Vital signs and toe-pinch response were monitored throughout the procedure and recorded every 10 minutes. The concentration of isoflurane was adjusted whenever necessary (e.g., slow or fast breathing), and the experiment was paused until stabilization of the values.

Once the physiological parameters stabilized, the shaved abdomen was sterilized with 3 alternating scrubs of 7.5% povidone-iodine (Betadine Surgical Scrub) and 70% alcohol, and sterile ultrasound gel (Sterile Aquasonic 100 gel) was applied. The sterile field was created, and thereafter, the only items that came into contact with the skin were the sterile guide and transducer. A critical aspect in this procedure is maintaining a straight approach to the target and avoiding the displacement of the fetus during penetration of the abdomen with the needle. Advancing a needle through the skin without displacing the underlying organs is challenging because of the toughness and elasticity of the dermal layer (Supplementary Vid. 1). In pilot experiments, we determined that the most reliable and efficient means to achieve precise targeting with minimal tissue displacement was to implement a three-step penetration procedure. In the first step, the guide tube is advanced through the ultrasound gel and aligned to the ultrasound transducer over the fetal head using the setup’s translational stages and stereotaxic arm manipulators. Depending on the positioning of individual fetuses, the angle of the guide tube is adjusted to accommodate the planned trajectory. While it gently presses against the skin, a brief electrical current is applied to the blunt non-insulated side of the guide tube to help insertion with minimal displacement of the fetus (Supplementary Vid. 2). As the injection needle can penetrate through the abdominal wall and into the uterus, insertion of the guide tube beyond the subcutaneous compartment is not necessary. This cautery step typically introduces air bubbles at the site of skin penetration. However, these bubbles cause minimal disruption of the US visualization of the target, since the angular approach means that the bubbles concentrate outside the US field of view. In the second step, the internal needle is advanced smoothly by hand through the guide, into the abdominal wall and uterine muscle and then into the lumen adjacent to the fetal skull. The position and angle of the needle relative to the head is important, with the desired approach angle determined, in part, by skull thickness and size of the uterine lumen, parameters that varied with species and gestational age. In the third step, the skull is manually penetrated with a brief jolt that abruptly displaces the needle tip by 1-3 mm (Supplementary Vid. 3). Penetration is most successful when attempted from an angle having maximum mechanical advantage, minimizing the risk of slipping along the convex skull surface and in a region of the cranium with thinner developing skull. Each attempt at penetration is evaluated in real time by the experimenter on the US display. In the case of trajectory misalignment or failed penetration, it is usually possible to readjust the geometrical approach to achieve successful entry without removal of the needle from the uterine lumen. A particularly effective strategy for this is to move the animal’s position and angle using the adjustments in the cradle setup with the guide tube still in place. Supplementary Vid. 3 shows a P107 FIVI in marmoset.

The rAAV virus, stored at -80°C, was transported to the procedure room on wet ice. After allowing it to thaw, the vector was thoroughly mixed using a micropipette, and the amount to inject was aliquoted onto a sterile syringe cap and loaded into the injection syringe. In the rat we injected between 2 and 10μL (eye socket: 2 μL; cerebral ventricles: 5 and 10 μL) and in the marmoset 10 and 60μL. The presence of small air bubbles in the last portion of the injection served to increase the echogenicity of the injectate and thus allowed visualization and confirmation of the injected target. Following an injection, the backflow of rAAV was reduced by keeping the needle in place for up to 5 minutes. Upon retraction of the needle and the guide tube, the heart of the fetus was monitored to ensure proper functioning and stability. These steps were repeated for each fetus. When determining the best path for an injection, special care was taken to avoid injuring nipples (in rats), internal organs (such as bladder and intestines) and large vessels (such as the umbilical cord). Whenever possible, we avoided penetrating the placenta to reach the fetus by repositioning the pregnant dam using the geared head and translational plates. On occasion, particularly in marmosets, injections were made through the placenta. When doing so, we used the color doppler mode to minimize the risks of and evaluate any bleeding along the trajectory. For most injection sites, no blood was observed upon removal of the needle and needle guide. On rare cases, a minimal amount was seen when retracting the needle, without any presence of active bleeding. At the end of the procedure, a small amount of local anesthetic medication (lidocaine) was applied to each site to prevent pain, along with a triple antibiotic ointment to prevent infection. Before returning to the housing room, the pregnant dam recovered in a warmed environment and was only transferred after being bright, alert, and reactive. The use of current to insert the guide resulted in mild scab formation on the injection sites in the days following the procedure. Typically, within one to two weeks, the scabs resolved without infections nor other complications. After each procedure, the pregnant dam was provided with supplemental nutritional support and monitored for any signs of pain, miscarriage (e.g., blood discharge, contractions) and dystocia until delivery. Supplementary table 3 provides additional information on the tools and agents used.

### Perinatal identification of positive animals

Being able to identify the progeny expressing a given transgene in an easy and timely manner is important to select animals for future experiments. For instance, the litters of rats injected in this procedure had up to 17 animals, and of those not all received FIVI injections. The high yield of endogenous reporter expression leads to brightness levels that can be observed using whole body imaging in living animals using LED flashlights. As for transgenic animals^5^, this approach was also effective in retrieving animals transduced in utero with viral vectors leading to strong expression of the reporter gene. In rats, expression was most prominent in the brain, spinal cord, back muscle and in the face (peripheral nerve labeling). In marmosets, it could be observed in less pigmented and hairless regions. Since reporter proteins are spectrally distinct, their distribution can be detected using LED flashlights with appropriate excitation and emission filters. We used a portable, battery-operated dual fluorescence flashlight with high intensity LEDs and Royal Blue and Green excitation filters (Model DFP-1, Nightsea™). For green shifted reporters we used the Royal Blue filter set, with excitation 440-460nm and 500-560nm bandpass barrier filter glasses. For red shifted reporters, we used the Green filter set, with excitation 510-540nm and 600nm bandpass barrier filter glasses. Transgene expression remains strong through adulthood, remaining visible in hairless regions.

### Histological processing

#### Perfusion and tissue collection

A 2-3 day variation in gestation is expected for the Sprague-Dawley SD strain. For this reason, to minimize the potential impact of age in the transduction patterns observed, the date of tissue collection in rats was determined based on postconceptional (PC) days rather than relative to birth. Unless otherwise indicated, all tissue collected from rats was obtained at PC44. At this age, approximately 21 days after birth, the number of neuronal cells in the rat brain, particularly in the cerebral cortex, is stable and maintained throughout adulthood^6^. Rats were deeply anesthetized with isoflurane gas (3-4% induction, 4-5% maintenance), and transcardially perfused with heparinized normal saline or 1X PBS followed by 4% paraformaldehyde (PFA, pH 7.4) in 1X phosphate-buffered saline (PBS). Marmosets were first deeply anesthetized with isoflurane gas (3-5%), and were overdosed by intraperitoneal (i.p.) delivery of pentobarbital (at least 80-120 mg/kg). They were then transcardially perfused as described above. In both rats and marmosets, the brains were extracted and postfixed in 4% PFA in 1X PBS. Note that the histological data presented in this study were obtained from animals injected with 10µL in rats and 60µL in marmosets.

#### Tissue selection and processing

In the most recent cases, following brain extraction, images of the dorsal and ventral surfaces of the fixed brains were acquired with a customized MacroFluo Z6 APO with a 0.5x Planapo Z-series objective (nA max. 0.0585), a digital monochrome DFC3000G camera, a LED3000 RL ring light, a fluorescence illuminator LRF 4/22 and external light source EL6000 allowing for both brightfield and fluorescence imaging (Leica Microsystems Inc.). The pictures were then exported to TIF format using the Leica Application Suite X, loaded into stack into Adobe Photoshop, and compared. In general, the brains with higher expression levels were selected for further processing, and the remaining were transferred to 1X PBS with 0.02% of sodium azide at 4°C for longer term storage. Some examples of these images are shown in Extended Data Figure 2. For the experiments in which gene expression was sparse, such as those with CrispR/Cas9, samples of spinal cords were sectioned to confirm expression. The brains selected for further processing were then cryopreserved in gradients of glycerol or sucrose (10 to 20 or 30%) in 1X PBS. The postfixation and cryoprotection intervals were adjusted based on the quality of the fixation and cryoprotection. Once cryoprotected, brains were sometimes embedded in 15% gelatin in 1X PBS. Forty micrometers thick sections were cut on a sliding freezing microtome or cryostat. One in twelve sections were collected in 1X PBS for screening, and the remaining sections were collected in an antifreeze solution for long-term storage and subsequent processing.

#### Immunofluorescent and microscopy imaging

Forty-micron thick free-floating sections were rinsed at room temperature (RT) with 1X PBS. Six 10-minute washes were performed for antifreeze-stored sections, and three washes for sections collected in 1XPBS. To permeabilize the cells, the sections were incubated in three 10-minute washes of 0.5% Triton X-100, followed by 30 minutes in 50% ethanol. After three 10-minute washes in 1X PBS, the sections were incubated in 10% Normal Goat Serum for 1 hour to block non-specific sites. They were then transferred to a PBST (1X PBS plus 0.05% Triton X-100) solution containing the primary antibody and incubated overnight at 4°C under gentle shaking. Supplementary Table 4 provides additional detail for the primary and secondary antibodies used in this study, including their working dilutions. For the marmoset fetal sections and rat CrispR/Cas9 and retinofugal projections cases, the endogenous reporter gene expression was enhanced with an anti-GFP antibody. The sections were then rinsed three times in 1X PBS and incubated in Alexa-fluor conjugated fluorescent secondary antibodies diluted with 1X PBS with gentle shaking at RT for 2h. Following three 10-minute rinses in 1X PBS, the sections were mounted in charged microscope slides, stained with Dapi and coverslipped with Fluoromount-G Mounting Medium to minimize fluorescence quenching and photobleaching during microscopy imaging. Sections were imaged through 20X objectives on a VS200 slide scanner from Olympus™ to obtain widefield images. Tiled z-stacks of selected regions were taken on a Leica Stellaris 8 confocal microscope using 20X or 40X objectives. Olympus, Leica or FIJI software were used to export the images after exposure adjustments and contrast enhancement, and to perform background subtraction when needed. When necessary, further adjustments in orientation and in brightness and contrast were made in Photoshop. For the PC87 case in Fig. 2, Photoshop was used to correct overexposure in the hemisphere with the brightest expression, allowing for improved visualization of the spatial symmetry of transduction.

#### Optical clearing and imaging of fetuses

A procedure based on the SHIELD^7^ (SHIELD fixation kit from Life Canvas, Inc) and CUBIC methods^8^ (reagents from Sigma-Aldrich, based on recipes in the cited manuscript) was used to evaluate labeling of the peripheral nervous system in intact fetuses. The fetus in Fig. 5e was co-injected with AAV9-CAG-tdTomato (5µl) and AAV2-CAG-GFP (5µl) at PC15. Only the AAV9-CAG-tdTomato is shown, as it is the only one labeling the peripheral nervous system with injections at this gestation time. The pregnant dam was transcardially perfused with heparinized normal saline, followed by 200mL of SHIELD fixation solution, at PC19, four days after the injections. After perfusion, positive fetuses were collected and postfixed in SHIELD fixation solution for 6 days, incubated in SHIELD OFF solution for 6 days, and transferred to 37°C SHIELD ON solution for 24 hours. Subsequently, they were transferred to 1X PBS with 0.1% sodium azide at 4°C until processing. The fetuses were passively cleared with Sodium dodecyl sulfate (SDS) clearing solution. Once cleared, they were transferred to 50% CUBIC-R+(N) overnight, and then 100% CUBIC-R+(N) for refractive index matching. Image acquisition was performed in silicone fluid (PM125, Clearco Products) using a 3i CTLS lightsheet microscope (1X, 0.25 NA objective, optical zoom 3.7). Image fields were merged using 3i Slidebook software and then background subtraction was performed using custom python scripts (adapted from de Rooi and colleages^9^) on the NIH High Performance Cluster, followed by importing the images into Arivis Vision4D for visualization by adjusting orientation and look up tables.

#### Quantification of NeuN and GFP neuronal co-labeling

Confocal images obtained in the Leica Stellaris 8 confocal microscope with a 20X objective were quantified using ImageJ software. Each image was smoothened with a 1µm median filter. Images were thresholded based on visual impression of the cell bodies and then binarized. Particle finder feature was used to filter and quantify blobs in the size range of 49-800µm. Each image was manually examined for blobs that contained multiple cells or non-cells (artifacts) and the final counts were adjusted. After applying this process to both GFP and NeuN channels, the images were merged, and the percentage of NeuN cells co-labeled with GFP cells was computed. The great majority of the GFP cells were NeuN-positive; only a minimal number were NeuN-negative.

---

**Extended Data Fig. 1:**
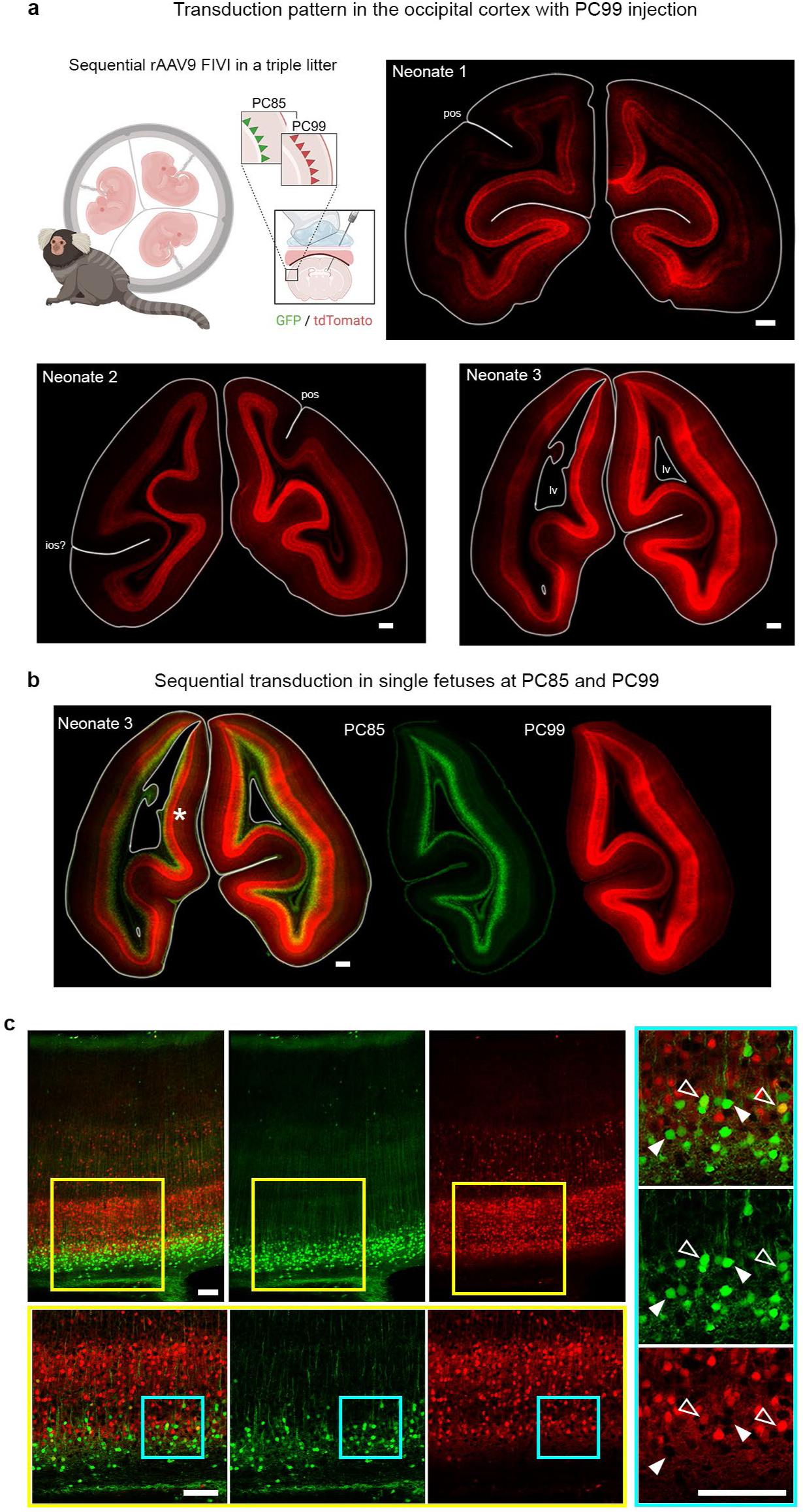
Sequential rAAV9 FIVI with distinct fluorescence reporters enable temporal and spatial mapping of cortical transduction. **a**, representative coronal sections from the occipital pole of a triplet litter injected sequentially with the same viral construct, rAAV9-CAG, carrying two different fluorescence reporters. A similar pattern of dense and widespread labeling for the FIVI at PC99 is observed perinatally in the lower cortical layers for all three neonates. **b**, representative section showing the labeling pattern from FIVI at PC85 and 99 in neonate 3, both generating continuous labeling across the cortex. **c**, comparative analysis of transduction patterns in neonate 3 from early and late injections reveals complementary labeling across cortical territories. The insets were selected from the midline cortex (region marked by an asterisk). While the labeled depths overlap, the cells transduced at each gestation timepoint are distinct (e.g., arrowheads showing GFP positive cells located within dense neuropil from tdTomato positive neurons), with occurrences of double-transduced neurons (open arrowhead) in the region of overlap of both populations. The extended gestation period of marmosets (∼143 days) enables sequential injections at intervals of days or months, providing the temporal resolution necessary to investigate spatial and temporal differences in cortical transduction with FIVI at different gestation times. Figure panel partially created in BioRender. Scale bars, a and b: 500 µm; c: 100 µm. Tissue collection: Neonate 1 at P10; Neonate 2 and 3 at P2. FIVI injection volumes at each timepoint: 60µL.

**Extended Data Fig. 2:**
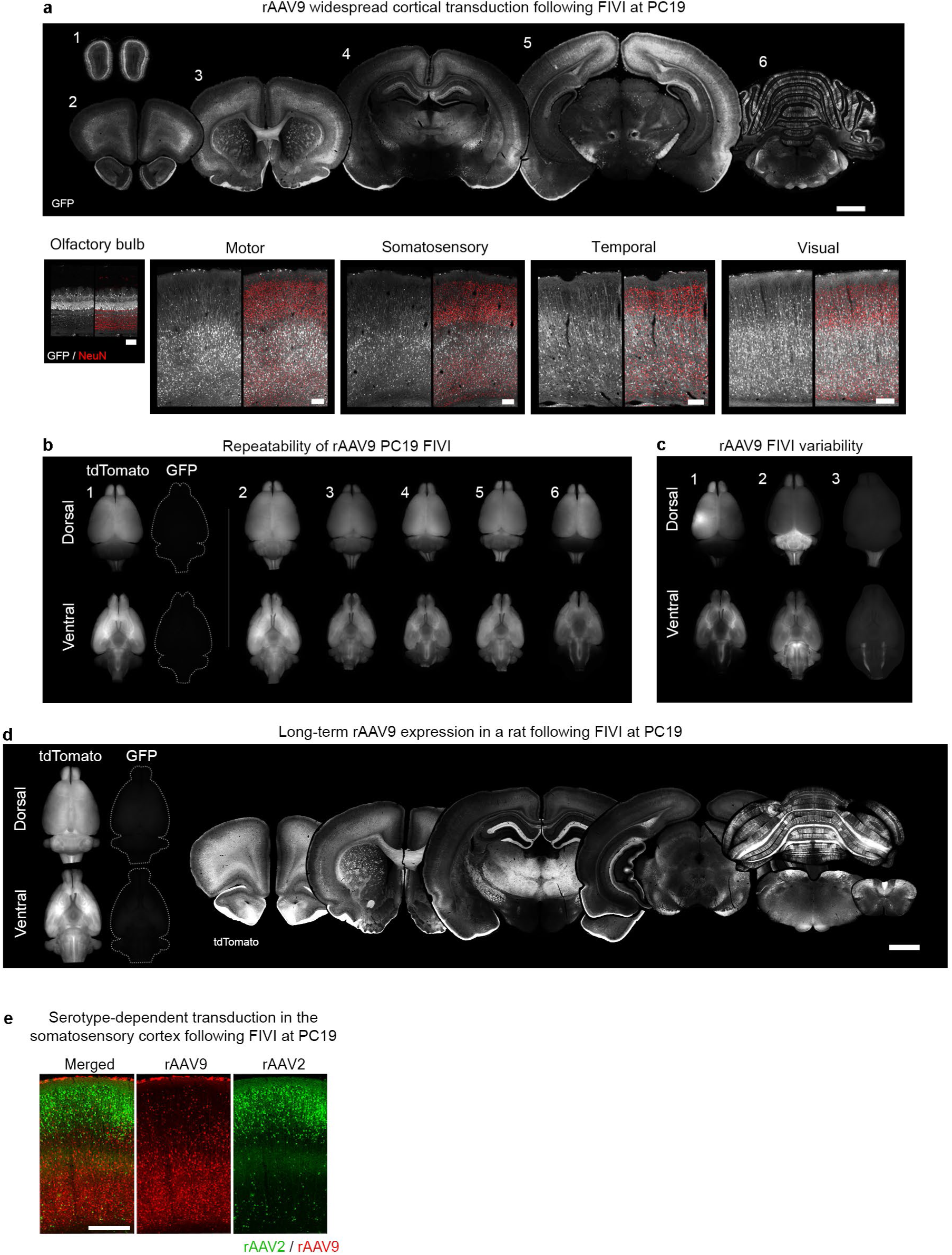
rAAV transduction patterns in rats following FIVI at PC19. **a**, exemplar coronal sections of a PC44 rat following FIVI at PC19 with rAAV9-CAG-GFP. **b**, dorsal and ventral views of PC44 brains injected at PC19 with rAAV9-CAG-tdTomato, collected at PC44. For brain 1, tdTomato expression is compared to background GFP expression under the same exposure parameters. Note the widespread transduction of the cortex and the repeatability of the cortical pattern across brains. **c**, examples of occasional variable labeling patterns at PC44 following FIVI at PC19. These include higher transduction within the presumptive injection tract in brain 1, and preferential transduction of posterior structures in brains 2 and 3 likely due to greater rAAV diffusion in the aqueduct and spinal canal. **d**, Long-term expression in the adult rat brain (5 months postnatal), injected in utero at PC19 with rAAV9-CAG-tdTomato, is shown in the dorsal and ventral surfaces as well as coronal sections. Note the strong reporter gene expression across the brain. **e**, exemplar transduction of the somatosensory cortex of a PC19 rat, displaying the differential patterns for rAAV2 and rAAV9. rAAV2 predominantly targets cells in more superficial layers, while rAAV9 targets cells deeper to AAV2-positive cells. Scale bars, 1 mm. FIVI injection volumes: 10µL.

**Extended Data Fig. 3:**
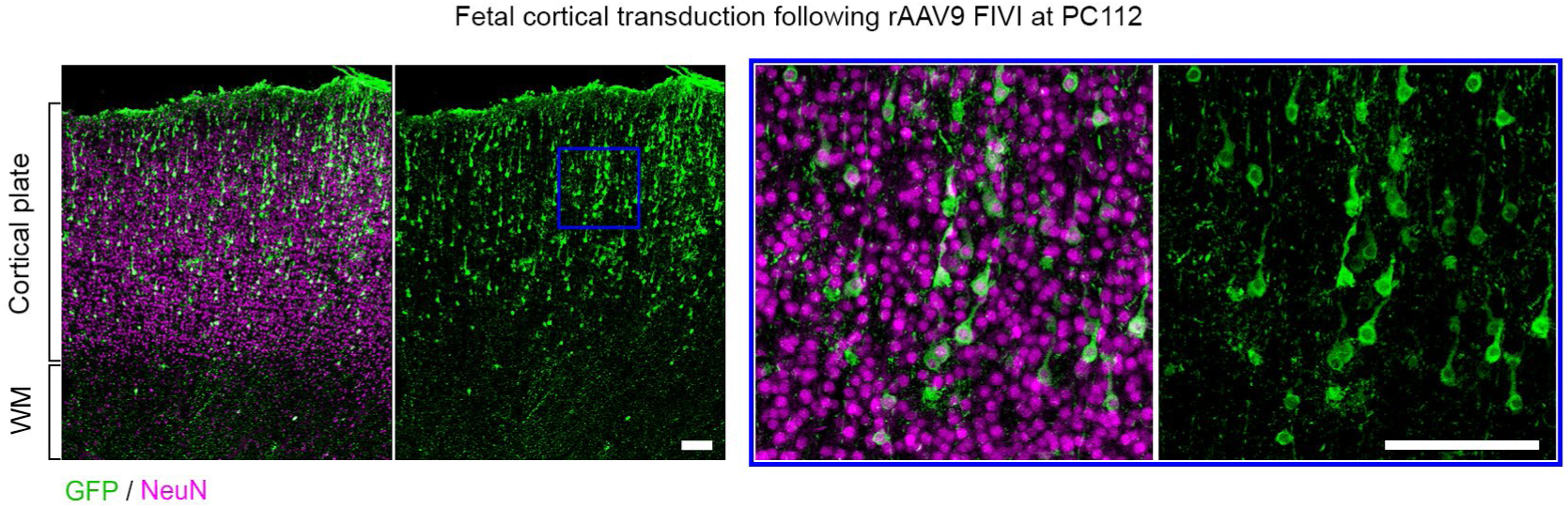
Transduction in the fetal marmoset cortex following FIVI at PC112. Prenatal transduction of pyramidal neurons in the cingulate cortex of a PC120 marmoset fetus injected in utero at PC112 with of rAAV9-CAG-GFP, showing labeling in more superficial layers consistent with the later gestational timing of the injection. Note that the tissue was collected following a miscarriage, resulting in limited analysis. Scale bars, 100 µm. FIVI injection volume: 60µL.

**Extended Data Fig. 4:**
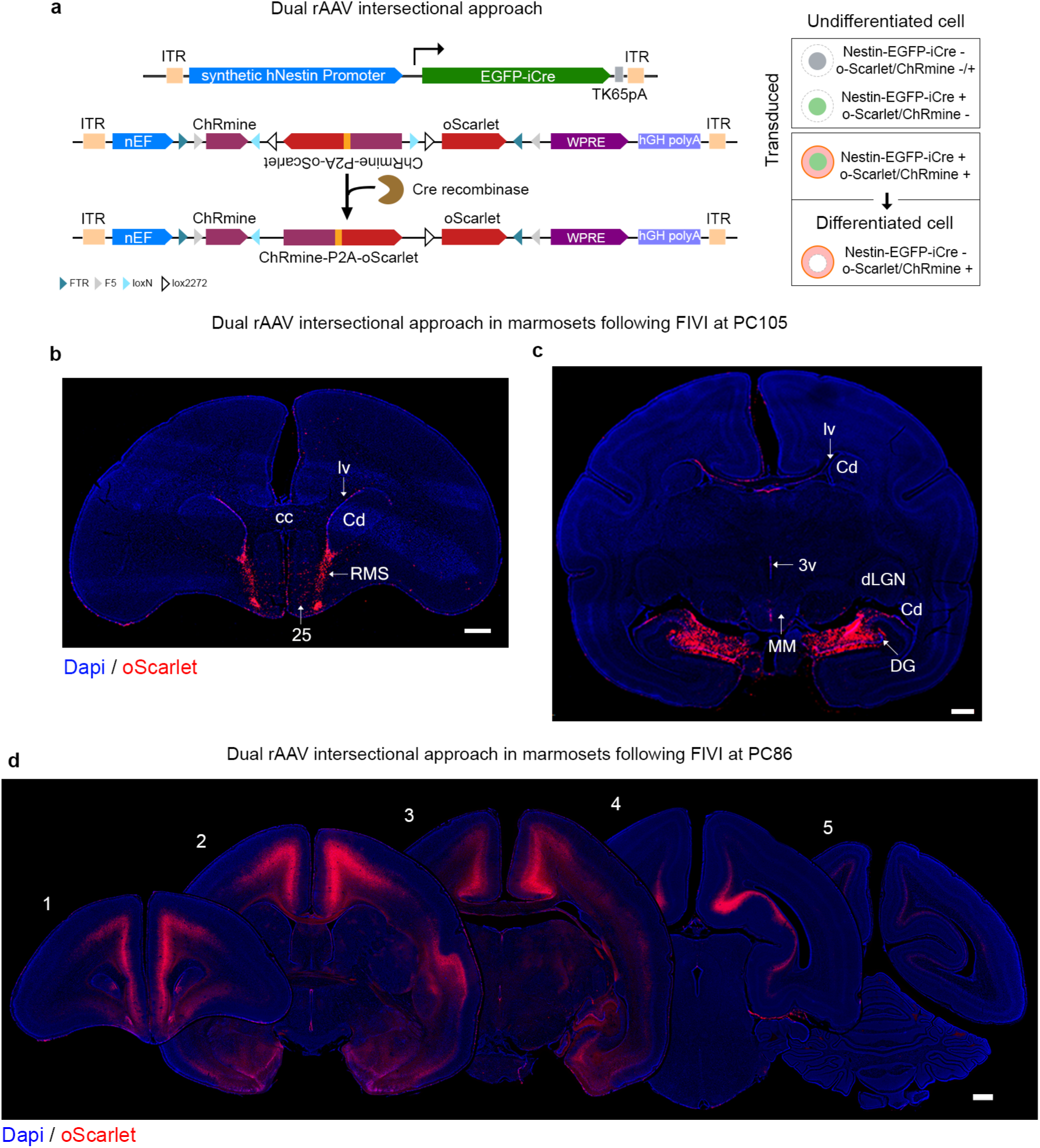
Intersectional genetic modification of the marmoset brain using cell-selective Cre expression to isolate Nestin-expressing cells during prenatal development. **a**, rAAV transduction is conditionally induced in Nestin-expressing cells through FIVI. Upon expression of Nestin, Cre recombinase is produced and mediates the inversion of “exon2” in the co-injected AAV8-nEF-Con/Foff 2.0-ChRmine-oScarlet construct, restoring ChRmine and oScarlet into the correct reading frame, leading to their expression under the constitutive nEF promoter. Singly transduced cells with either construct or cells not expressing Nestin will not produce ChRmine and oScarlet. Only the cells transduced with both constructs and expressing Nestin after the injection will express these transgenes. Cre recombinase mediates the permanent inversion of transgenes into the correct reading frame, allowing for their continued expression even after the cells differentiate and stop expressing Nestin. This strategy is effective for evaluating which cells had transient expression of a gene of interest at a specific time. **b** and **c** show selective labeling, bilaterally, in a postnatal day 5 marmoset in regions suggested to be involved in late and postnatal neurogenesis and maturation, following a PC105 FIVI co-injection of the constructs in **a**. **b**, oScarlet+ expression can be seen in cells extending from the lateral ventricle (lv) to the olfactory regions, following the path of the rostral migratory stream (RMS). Additional cells extend to the midline (area 25), suggestive of the medial migratory stream. **c**. shows prominent labeling in the dentate gyrus of the hippocampus. Across the brain, there is dense labeling observed in the walls of the ventricles. **d**, shows reported expression following co-injection of these constructs at PC86. Compared to the injection performed at PC105, the earlier gestational timing resulted in broader cortical labeling, with transduced cells at postnatal day 2 predominantly located in the lower cortical layers. Scale bars, 1 mm. Nomenclature: 25: areas 25; 3V: 3^rd^ ventricle; Cd: caudate nucleus; cc: corpus callosum; DG: dentate gyrus; dLGN: lateral geniculate nucleus; lv: lateral ventricle; MM: medial mammillary bodies. FIVI injection volumes: 60µL.

## References

1 Luo, L., Callaway, E. M. & Svoboda, K. Genetic Dissection of Neural Circuits: A Decade of Progress. Neuron 98, 256–281 (2018). 10.1016/j.neuron.2018.03.040

2 Huang, Z. J. & Zeng, H. Genetic approaches to neural circuits in the mouse. Annual review of neuroscience 36, 183–215 (2013). 10.1146/annurev-neuro-062012-170307

3 Fenno, L., Yizhar, O. & Deisseroth, K. The development and application of optogenetics. Annual review of neuroscience 34, 389–412 (2011). 10.1146/annurev-neuro-061010-113817

4 Madisen, L. et al. A robust and high-throughput Cre reporting and characterization system for the whole mouse brain. Nat Neurosci 13, 133–140 (2010). 10.1038/nn.2467

5 Charreau, B., Tesson, L., Soulillou, J. P., Pourcel, C. & Anegon, I. Transgenesis in rats: technical aspects and models. Transgenic Res 5, 223–234 (1996). 10.1007/BF01972876

6 Chenouard, V. et al. Advances in Genome Editing and Application to the Generation of Genetically Modified Rat Models. Front Genet 12, 615491 (2021). 10.3389/fgene.2021.615491

7 Johnson, M. B. et al. Aspm knockout ferret reveals an evolutionary mechanism governing cerebral cortical size. Nature 556, 370–375 (2018). 10.1038/s41586-018-0035-0

8 Yu, M. et al. Highly Efficient Transgenesis in Ferrets Using CRISPR/Cas9-Mediated Homology-Independent Insertion at the ROSA26 Locus. Sci Rep 9, 1971 (2019). 10.1038/s41598-018-37192-4

9 Park, J. E. et al. Generation of transgenic marmosets expressing genetically encoded calcium indicators. Sci Rep 6, 34931 (2016). 10.1038/srep34931

10 Sasaki, E. et al. Generation of transgenic non-human primates with germline transmission. Nature 459, 523–527 (2009). 10.1038/nature08090

11 Yang, S. H. et al. Towards a transgenic model of Huntington’s disease in a non-human primate. Nature 453, 921–924 (2008). 10.1038/nature06975

12 Niu, Y. et al. Generation of gene-modified cynomolgus monkey via Cas9/RNA-mediated gene targeting in one-cell embryos. Cell 156, 836–843 (2014). 10.1016/j.cell.2014.01.027

13 Liu, Z. et al. Autism-like behaviours and germline transmission in transgenic monkeys overexpressing MeCP2. Nature 530, 98–102 (2016). 10.1038/nature16533

14 Izpisua Belmonte, J. C., et al. Brains, genes, and primates. Neuron 86, 617–631 (2015). 10.1016/j.neuron.2015.03.021

15 Emborg, M. E. Nonhuman Primate Models of Neurodegenerative Disorders. ILAR J 58, 190–201 (2017). 10.1093/ilar/ilx021

16 Jennings, C. G. et al. Opportunities and challenges in modeling human brain disorders in transgenic primates. Nat Neurosci 19, 1123–1130 (2016). 10.1038/nn.4362

17 Feng, G. et al. Opportunities and limitations of genetically modified nonhuman primate models for neuroscience research. Proceedings of the National Academy of Sciences of the United States of America 117, 24022–24031 (2020). 10.1073/pnas.2006515117

18 Aida, T. & Feng, G. The dawn of non-human primate models for neurodevelopmental disorders. Curr Opin Genet Dev 65, 160–168 (2020). 10.1016/j.gde.2020.05.040

19 Nectow, A. R. & Nestler, E. J. Viral tools for neuroscience. Nat Rev Neurosci 21, 669–681 (2020). 10.1038/s41583-020-00382-z

20 Campos, L. J. et al. Advances in AAV technology for delivering genetically encoded cargo to the nonhuman primate nervous system. Curr Res Neurobiol 4, 100086 (2023). 10.1016/j.crneur.2023.100086

21 Kim, J. Y. et al. Viral transduction of the neonatal brain delivers controllable genetic mosaicism for visualising and manipulating neuronal circuits in vivo. Eur J Neurosci 37, 1203–1220 (2013). 10.1111/ejn.12126

22 Clancy, B., Darlington, R. B. & Finlay, B. L. Translating developmental time across mammalian species. Neuroscience 105, 7–17 (2001). 10.1016/s0306-4522(01)00171-3

23 Rahim, A. A. et al. In utero administration of Ad5 and AAV pseudotypes to the fetal brain leads to efficient, widespread and long-term gene expression. Gene Ther 19, 936–946 (2012). 10.1038/gt.2011.157

24 Massaro, G. et al. Fetal gene therapy for neurodegenerative disease of infants. Nature medicine 24, 1317–1323 (2018). 10.1038/s41591-018-0106-7

25 Charvet, C. J. & Finlay, B. L. Evo-devo and the primate isocortex: the central organizing role of intrinsic gradients of neurogenesis. Brain Behav Evol 84, 81–92 (2014). 10.1159/000365181

26 Kolk, S. M. & Rakic, P. Development of prefrontal cortex. Neuropsychopharmacology 47, 41–57 (2022). 10.1038/s41386-021-01137-9

27 Rakic, P. Neurons in rhesus monkey visual cortex: systematic relation between time of origin and eventual disposition. Science 183, 425–427 (1974). 10.1126/science.183.4123.425

28 Rakic, P. Pre- and post-developmental neurogenesis in primates. Clinical Neuroscience Research 2 (2002). 10.1016/S1566-2772(02)00005-1

29 Rakic, P. Specification of cerebral cortical areas. Science 241, 170–176 (1988). 10.1126/science.3291116

30 Meyer, N. L. & Chapman, M. S. Adeno-associated virus (AAV) cell entry: structural insights. Trends Microbiol 30, 432–451 (2022). 10.1016/j.tim.2021.09.005

31 Di Bella, D. J., Dominguez-Iturza, N., Brown, J. R. & Arlotta, P. Making Ramon y Cajal proud: Development of cell identity and diversity in the cerebral cortex. Neuron 112, 2091–2111 (2024). 10.1016/j.neuron.2024.04.021

32 Bernal, A. & Arranz, L. Nestin-expressing progenitor cells: function, identity and therapeutic implications. Cell Mol Life Sci 75, 2177–2195 (2018). 10.1007/s00018-018-2794-z

33 Bott, C. J. et al. Nestin in immature embryonic neurons affects axon growth cone morphology and Semaphorin3a sensitivity. Mol Biol Cell 30, 1214–1229 (2019). 10.1091/mbc.E18-06-0361

34 Noisa, P., Urrutikoetxea-Uriguen, A., Li, M. & Cui, W. Generation of human embryonic stem cell reporter lines expressing GFP specifically in neural progenitors. Stem Cell Rev Rep 6, 438–449 (2010). 10.1007/s12015-010-9159-9

35 Doudna, J. A. The promise and challenge of therapeutic genome editing. Nature 578, 229–236 (2020). 10.1038/s41586-020-1978-5

36 Nishiyama, J., Mikuni, T. & Yasuda, R. Virus-Mediated Genome Editing via Homology-Directed Repair in Mitotic and Postmitotic Cells in Mammalian Brain. Neuron 96, 755–768 e755 (2017). 10.1016/j.neuron.2017.10.004

37 Ayers, J. I. et al. Widespread and efficient transduction of spinal cord and brain following neonatal AAV injection and potential disease modifying effect in ALS mice. Mol Ther 23, 53–62 (2015). 10.1038/mt.2014.180

38 Tremblay, S. et al. An Open Resource for Non-human Primate Optogenetics. Neuron 108, 1075–1090 e1076 (2020). 10.1016/j.neuron.2020.09.027

39 Mattar, C. N. et al. Systemic delivery of scAAV9 in fetal macaques facilitates neuronal transduction of the central and peripheral nervous systems. Gene Ther 20, 69–83 (2013). 10.1038/gt.2011.216

40 Dehay, B., Dalkara, D., Dovero, S., Li, Q. & Bezard, E. Systemic scAAV9 variant mediates brain transduction in newborn rhesus macaques. Sci Rep 2, 253 (2012). 10.1038/srep00253

41 Kim, S. et al. Whole-brain mapping of effective connectivity by fMRI with cortex-wide patterned optogenetics. Neuron 111, 1732–1747 e1736 (2023). 10.1016/j.neuron.2023.03.002

42 Song, X. et al. Mesoscopic landscape of cortical functions revealed by through-skull wide-field optical imaging in marmoset monkeys. Nat Commun 13, 2238 (2022). 10.1038/s41467-022-29864-7

43 Musall, S., Kaufman, M. T., Juavinett, A. L., Gluf, S. & Churchland, A. K. Single-trial neural dynamics are dominated by richly varied movements. Nat Neurosci 22, 1677–1686 (2019). 10.1038/s41593-019-0502-4

44 Fredericks, J. M. et al. Methods for mechanical delivery of viral vectors into rhesus monkey brain. J Neurosci Methods 339, 108730 (2020). 10.1016/j.jneumeth.2020.108730

45 Lerchner, W. et al. Efficient viral expression of a chemogenetic receptor in the old-world monkey amygdala. Curr Res Neurobiol 4, 100091 (2023). 10.1016/j.crneur.2023.100091

46 Yazdan-Shahmorad, A. et al. Widespread optogenetic expression in macaque cortex obtained with MR-guided, convection enhanced delivery (CED) of AAV vector to the thalamus. J Neurosci Methods 293, 347–358 (2018). 10.1016/j.jneumeth.2017.10.009

47 Foust, K. D. et al. Intravascular AAV9 preferentially targets neonatal neurons and adult astrocytes. Nat Biotechnol 27, 59–65 (2009). 10.1038/nbt.1515

48 Gray, S. J. et al. Preclinical differences of intravascular AAV9 delivery to neurons and glia: a comparative study of adult mice and nonhuman primates. Mol Ther 19, 1058–1069 (2011). 10.1038/mt.2011.72

49 Samaranch, L. et al. Adeno-associated virus serotype 9 transduction in the central nervous system of nonhuman primates. Hum Gene Ther 23, 382–389 (2012). 10.1089/hum.2011.200

50 Chan, K. Y. et al. Engineered AAVs for efficient noninvasive gene delivery to the central and peripheral nervous systems. Nat Neurosci 20, 1172–1179 (2017). 10.1038/nn.4593

51 Deverman, B. E. et al. Cre-dependent selection yields AAV variants for widespread gene transfer to the adult brain. Nat Biotechnol 34, 204–209 (2016). 10.1038/nbt.3440

52 Bedbrook, C. N., Deverman, B. E. & Gradinaru, V. Viral Strategies for Targeting the Central and Peripheral Nervous Systems. Annual review of neuroscience 41, 323–348 (2018). 10.1146/annurev-neuro-080317-062048

53 Chuapoco, M. R. et al. Adeno-associated viral vectors for functional intravenous gene transfer throughout the non-human primate brain. Nat Nanotechnol (2023). 10.1038/s41565-023-01419-x

54 Goertsen, D. et al. AAV capsid variants with brain-wide transgene expression and decreased liver targeting after intravenous delivery in mouse and marmoset. Nat Neurosci 25, 106–115 (2022). 10.1038/s41593-021-00969-4

55 Hammond, S. L., Leek, A. N., Richman, E. H. & Tjalkens, R. B. Cellular selectivity of AAV serotypes for gene delivery in neurons and astrocytes by neonatal intracerebroventricular injection. PloS one 12, e0188830 (2017). 10.1371/journal.pone.0188830

56 Tabata, H. & Nakajima, K. Efficient in utero gene transfer system to the developing mouse brain using electroporation: visualization of neuronal migration in the developing cortex. Neuroscience 103, 865–872 (2001). 10.1016/s0306-4522(01)00016-1

57 Taniguchi, Y., Young-Pearse, T., Sawa, A. & Kamiya, A. In utero electroporation as a tool for genetic manipulation in vivo to study psychiatric disorders: from genes to circuits and behaviors. The Neuroscientist : a review journal bringing neurobiology, neurology and psychiatry 18, 169–179 (2012). 10.1177/1073858411399925

58 Yamashiro, K., Ikegaya, Y. & Matsumoto, N. In Utero Electroporation for Manipulation of Specific Neuronal Populations. Membranes (Basel) 12 (2022). 10.3390/membranes12050513

59 Harrison, M. R., Anderson, J., Rosen, M. A., Ross, N. A. & Hendrickx, A. G. Fetal surgery in the primate I. Anesthetic, surgical, and tocolytic management to maximize fetal-neonatal survival. J Pediatr Surg 17, 115–122 (1982). 10.1016/s0022-3468(82)80193-0

60 Adzick, N. S. et al. Fetal surgery in the primate. III. Maternal outcome after fetal surgery. J Pediatr Surg 21, 477–480 (1986). 10.1016/s0022-3468(86)80215-9

61 Rakic, P. & Goldman-Rakic, P. S. in Perinatal Neurology and Neurosurgery (eds Richard A. Thompson, John R. Green, & Stanley D. Johnsen) 1–15 (Springer Netherlands, 1985).

62 An, M., et al. Engineered virus-like particles for transient delivery of prime editor ribonucleoprotein complexes in vivo. Nat Biotechnol 42, 1526–1537 (2024). 10.1038/s41587-023-02078-y

63 Hamilton, J. R. et al. In vivo human T cell engineering with enveloped delivery vehicles. Nat Biotechnol 42, 1684–1692 (2024). 10.1038/s41587-023-02085-z

64 Mallapaty, S. Rare genetic disorder treated in womb for the first time. Nature 638, 869 (2025). 10.1038/d41586-025-00534-0

65 Newby, G. A. et al. Base editing of haematopoietic stem cells rescues sickle cell disease in mice. Nature 595, 295–302 (2021). 10.1038/s41586-021-03609-w

66 Regev, L., Ezrielev, E., Gershon, E., Gil, S. & Chen, A. Genetic approach for intracerebroventricular delivery. Proceedings of the National Academy of Sciences of the United States of America 107, 4424–4429 (2010). 10.1073/pnas.0907059107

67 Jang, A. & Lehtinen, M. K. In utero intracerebroventricular delivery of adeno-associated viral vectors to target mouse choroid plexus and cerebrospinal fluid. STAR Protoc 4, 101975 (2023). 10.1016/j.xpro.2022.101975

68 Jang, A. & Lehtinen, M. K. Experimental approaches for manipulating choroid plexus epithelial cells. Fluids Barriers CNS 19, 36 (2022). 10.1186/s12987-022-00330-2

69 Chen, S. H. et al. Recombinant Viral Vectors as Neuroscience Tools. Curr Protoc Neurosci 87, e67 (2019). 10.1002/cpns.67

70 Matsuzaki, Y., Fukai, Y., Konno, A. & Hirai, H. Optimal different adeno-associated virus capsid/promoter combinations to target specific cell types in the common marmoset cerebral cortex. Mol Ther Methods Clin Dev 32, 101337 (2024). 10.1016/j.omtm.2024.101337

71 Watakabe, A. et al. Comparative analyses of adeno-associated viral vector serotypes 1, 2, 5, 8 and 9 in marmoset, mouse and macaque cerebral cortex. Neurosci Res 93, 144–157 (2015). 10.1016/j.neures.2014.09.002

72 Srivastava, A. In vivo tissue-tropism of adeno-associated viral vectors. Curr Opin Virol 21, 75–80 (2016). 10.1016/j.coviro.2016.08.003

73 Bandtlow, C. E. & Zimmermann, D. R. Proteoglycans in the developing brain: new conceptual insights for old proteins. Physiol Rev 80, 1267–1290 (2000). 10.1152/physrev.2000.80.4.1267

74 Lee, B. & An, H. J. Small but big leaps towards neuroglycomics: exploring N-glycome in the brain to advance the understanding of brain development and function. Neural Regen Res 19, 489–490 (2024). 10.4103/1673-5374.380887

75 Rutherford, J. N., deMartelly, V. A., Layne Colon, D. G., Ross, C. N. & Tardif, S. D. Developmental origins of pregnancy loss in the adult female common marmoset monkey (Callithrix jacchus). PloS one 9, e96845 (2014). 10.1371/journal.pone.0096845

76 Tardif, S. D. et al. Reproduction in captive common marmosets (Callithrix jacchus). Comp Med 53, 364–368 (2003).

77 Ose, T. et al. Anatomical variability, multi-modal coordinate systems, and precision targeting in the marmoset brain. Neuroimage 250, 118965 (2022). 10.1016/j.neuroimage.2022.118965

78 Kita, Y. et al. Cellular-resolution gene expression profiling in the neonatal marmoset brain reveals dynamic species- and region-specific differences. Proceedings of the National Academy of Sciences of the United States of America 118 (2021). 10.1073/pnas.2020125118

79 Shimogori, T. et al. Digital gene atlas of neonate common marmoset brain. Neurosci Res 128, 1–13 (2018). 10.1016/j.neures.2017.10.009

80 Kremer, L. P. M. et al. High throughput screening of novel AAV capsids identifies variants for transduction of adult NSCs within the subventricular zone. Mol Ther Methods Clin Dev 23, 33–50 (2021). 10.1016/j.omtm.2021.07.001

81 Brown, N., Song, L., Kollu, N. R. & Hirsch, M. L. Adeno-Associated Virus Vectors and Stem Cells: Friends or Foes? Hum Gene Ther 28, 450–463 (2017). 10.1089/hum.2017.038

## References

1 Oerke, A. K., Einspanier, A. & Hodges, J. K. Detection of pregnancy and monitoring patterns of uterine and fetal growth in the marmoset monkey (Callithrix jacchus) by real-time ultrasonography. Am J Primatol 36, 1–13 (1995). 10.1002/ajp.1350360102

2 Noisa, P., Urrutikoetxea-Uriguen, A., Li, M. & Cui, W. Generation of human embryonic stem cell reporter lines expressing GFP specifically in neural progenitors. Stem Cell Rev Rep 6, 438–449 (2010). 10.1007/s12015-010-9159-9

3 Nishiyama, J., Mikuni, T. & Yasuda, R. Virus-Mediated Genome Editing via Homology-Directed Repair in Mitotic and Postmitotic Cells in Mammalian Brain. Neuron 96, 755–768 e755 (2017). 10.1016/j.neuron.2017.10.004

4 Howard, D. B., Powers, K., Wang, Y. & Harvey, B. K. Tropism and toxicity of adeno-associated viral vector serotypes 1, 2, 5, 6, 7, 8, and 9 in rat neurons and glia in vitro. Virology 372, 24–34 (2008). 10.1016/j.virol.2007.10.007

5 Tyas, D. A., Pratt, T., Simpson, T. I., Mason, J. O. & Price, D. J. Identifying GFP-transgenic animals by flashlight. Biotechniques 34, 474–476 (2003). 10.2144/03343bm04

6 Bandeira, F., Lent, R. & Herculano-Houzel, S. Changing numbers of neuronal and non-neuronal cells underlie postnatal brain growth in the rat. Proceedings of the National Academy of Sciences of the United States of America 106, 14108–14113 (2009). 10.1073/pnas.0804650106

7 Park, Y. G. et al. Protection of tissue physicochemical properties using polyfunctional crosslinkers. Nat Biotechnol (2018). 10.1038/nbt.4281

8 Susaki, E. A. et al. Advanced CUBIC protocols for whole-brain and whole-body clearing and imaging. Nat Protoc 10, 1709–1727 (2015). 10.1038/nprot.2015.085

9 de Rooi, J. J., Devos, O., Sliwa, M., Ruckebusch, C. & Eilers, P. H. Mixture models for two-dimensional baseline correction, applied to artifact elimination in time-resolved spectroscopy. Anal Chim Acta 771, 7–13 (2013). 10.1016/j.aca.2013.02.007

