## Supplementary Tables for "Targeted gene transfer into developmentally defined cell populations of the primate brain"

**Supplementary Table 1: Viral constructs used in this study**

| Serotype | AAV Vector | Packaged by | Unique ID | Addgene Plasmid | Titer range (VG/mL) | Gift from | Used for |
|---|---|---|---|---|---|---|---|
| AAV2 | pAAV-CAG-GFP | Addgene | 37825-AAV2 | 37825 | 5.30E+12 - 1.5E+13 | Edward Boyden | Serotype evaluation |
| AAV8 | pAAV-nEF-Con/Foff 2.0-ChRmine-oScarlet | Addgene | 137161-AAV8 | 137161 | 2.10E+13 | Karl Deisseroth & INTRSECT 2.0 Project | Cre/Lox recombination-based intersectional strategy |
| AAV9 | pAAV-CAG-GFP | Addgene | 37825-AAV9 | 37825 | 2.60E+13 | Edward Boyden | Gestation timing and serotype evaluation |
| AAV9 | pAAV-CAG-tdTomato (codon diversified) | Addgene | 59462-AAV9 | 59462 | 2.3E+13 - 2.5E+13 | Edward Boyden | Gestation timing evaluation |
| AAV9 | pAAV-NestinTK-EGFP-iCre | NIDA GEVVC | AAV916 | 228443 | 4.29E+12 | (in house) | Cre/Lox recombination-based intersectional strategy |
| AAV9 | pAAV-rActin-EGFP-donor | NIDA GEVVC | AAV942 | 228444 | 1.96E+13 | (in house) | CRISPR/Cas9-mediated genome editing |
| AAV9 | AAV9-EFS-SpCas9 | NIDA GEVVC | AAV949 | 104588 | 1.40E+13 | (in house) | CRISPR/Cas9-mediated genome editing |

**Supplementary Table 2: Plasmids used in this study**

| Plasmid | Source | Catalog number | Addgene Plasmid | Gift from | Used for |
|---|---|---|---|---|---|
| pNestin-EGFP | Addgene | 38777 | 38777 | Wei Cui | Template for cloning, pOTTC2234 |
| pAAV-HDR-mEGFP-Actin | Addgene | 119870 | 119870 | Ryohei Yasuda | Template for cloning, pOTTC2360 |
| pAAV-EFS-SpCas9 | Addgene | 104588 | 104588 | Ryohei Yasuda | Packaged as vector in top section |
| pAAV CMV-IE eGFP-2A-iCre | NIDA GEVVC | pOTTC1031 | n/a | NIDA GEVVC | Template for cloning, pOTTC2234 |
| pAAV JeT ConFon HA-hM4D(Gi)-mCherry TK65pA | NIDA GEVVC | pOTTC2210 | n/a | NIDA GEVVC | Template for cloning, pOTTC2234 |
| pAAV Nestin GFP-iCre | NIDA GEVVC | pOTTC2234 | 228443 | this study | Packaged as vector in top section |
| pAAV rActin EGFP donor | NIDA GEVVC | pOTTC2360 | 228444 | this study | Packaged as vector in top section |
| pHelper | PENN Vector Core | n/a | n/a | PENN Vector Core, Perelman School of Medicine, University of Pennsylvania | AAV packaging plasmids, in trans |
| pAAV 2/9 | PENN Vector Core | n/a | n/a | PENN Vector Core, Perelman School of Medicine, University of Pennsylvania | AAV packaging plasmids, in trans |

**Supplementary Table 3: Setup and procedure tools**

***Anesthesia machine and monitoring systems***

| *Product* | *Name* | *Part #* | *Brand* | *Reusable* | *Custom-made* |
|---|---|---|---|---|---|
| Anesthesia machine | RC2 Rodent Circuit Controller | 922100 | VetEquip | Y | N |
| Induction box (rat and neonatal marmosets) | Induction Chamber, 7 liter | 941488 | VetEquip | Y | N |
| Conductive gas supply hose | Color coded, nylon-reinforced, 1/4" ID, conductive gas supply hose | 931503 | VetEquip | Y | N |
| Rat nose cone | 12mm Nosecone and 14mm Nosecone | 921612 and 921614 | VetEquip | Y | N |
| Marmoset nose cone | Anesthesia silicone mask, reusable, size 00, premature baby | MP02900 | Drager | Y | N |
| Infrared heater | NORMOTHERM™ INFRARED HEATER | n/a | Britz & Company | Y | N |
| Rat monitoring system | SomnoSuite® Low-Flow Anesthesia System | SS-01 | Kent Scientific Corporation | Y | N |
| Marmoset monitoring system | IntelliVue MX500 Patient Monitor | 866064 | Philips Medizin Systeme Böblingen GmbH | Y | N |
| Marmoset EKG leads | Micro NeoLead, AAMI radio lead | 989803183141 | Philips Medical Systems Hsg | Y | N |
| SpO2 Sensor | Nasal Alar SpO2 Sensor | 989803205381 | Philips Medical Systems Hsg | Y | N |
| Blood pressure cuff | NIBP Cuffs (Neonatal single-patient cuff size #1) | M1866B | Philips Medical Systems Hsg | Y | N |

***Electrosurgery unit***

| *Product* | *Name* | *Part #* | *Brand* | *Reusable* | *Custom-made* |
|---|---|---|---|---|---|
| Electrosurgery Unit | Symmetry Pro-120 Multi-Pupose Electrosurgical Generator | A1250S | Bovie | Y | N |
| Electrosurgical Pencil | Push Button Electrosurgical Pencil 50/Box | ESP1 | Bovie | Y | N |
| Reusable Grounding Cable | Reusable Grounding Cable | A1252C | Bovie | Y | N |
| Return Electrodes | Disposable Split Adult Return Electrodes | ESRE-1 | Bovie | N | N |
| Sterile US gel | Aquasonic 100 - Sterile Single Use - Overwrapped Foil Pouches 48 per box | PLI 01-01 | Aquasonic | N | N |

**Guides and needles**

| *Product* | *Name* | *Part #* | *Brand* | *Reusable* | *Custom-made* |
|---|---|---|---|---|---|
| Guide tubes | 23 gauge, Hubless Needle, custom length (2 in), point style 4, 12 DEG, 6/PK | 22023-01 | Hamilton Company | Y | Y |
| | 24 gauge, Hubless Needle, custom length (2 in), point style 4, 12 DEG, 6/PK | 22024-01 | Hamilton Company | | |
| Needles | 33 gauge, Small Hub RN Needle, custom length (2.5 in), point style 4, 12 DEG, 6/PK | 7803-05 | Hamilton Company | Y | Y |
| | 31 gauge, Small Hub RN Needle, custom length (2.5 in), point style 4, 12 DEG, 6/PK | 7803-03 | Hamilton Company | | |
| Syringe | 5 µL, Model 75 RN Syringe, Needle Sold Separately | 7634-01 | Hamilton Company | Y | N |
| | 10 µL, Model 701 RN Syringe, Needle Sold Separately | 7635-01 | Hamilton Company | | |
| | 100 µL, Model 710 RN Syringe, Needle Sold Separately | 7638-01 | Hamilton Company | | |
| Heat shrink tubing | Palladium™ "Pebax™" Heat Shrink Tubing | PBST2-040-40-004C | Component Supply | N | N |
| Heat gun | 1500 Watt 10 Amp 12 Temperature Heat Gun | 69343 | Harborfreight | Y | N |

***Injection setup***

| *Product* | *Name* | *Part #* | *Brand* | *Reusable* | *Custom-made* |
|---|---|---|---|---|---|
| 18.70 mm A/P Bar | Model 1400 series | n/a | KOPF | Y | N |
| A.P. Slide Attachment | Model 1261 A/P Slide Attachment | n/a | KOPF | Y | N |
| Rotation Adapter | Model 1460-G Rotation Adapter | n/a | KOPF | Y | N |
| Travel Vertical Translation Stage | VAP4/M - 101.6 mm Travel Vertical Translation Stage, M4 and M6 Taps | VAP4/M | KOPF | Y | N |
| Stereotaxic arm and micro manipulator with fine adjustm | Model 1460-61 electrode Carrier with Fine Adjustment A/P Slide Assembly | 1460-61 | KOPF | Y | N |
| Single Axis Translation Stage | LT1/M - Single Axis Translation Stage, 50 mm Travel, Metric | LT1/M | THORLABS | Y | N |
| Cube Geared Head | Arca-Swiss C1 Cube Geared Head with Arca Classic Quick | 8501303.1 | Arca-Swiss | Y | N |
| Aluminum Breadboard | Aluminum Breadboard, 250 mm x 300 mm x 12.7 mm, M6 Taps | MB2530/M | THORLABS | Y | N |
| Animal cradle | n/a | n/a | NIMH/SI | Y | Y |
| Stereotaxic arm base holder | n/a | n/a | NIMH/SI | Y | Y |
| Ultrasound probe holder | n/a | n/a | NIMH/SI | Y | Y |
| Costum plates | n/a | n/a | NIMH/SI | Y | Y |
| Stainless steel extended spring clip | n/a | n/a | | Y | N |
| Loc-Line® 0.5" | n/a | n/a | | Y | N |

***Ultrasound machine and transducers***

| *Product* | *Name* | *Part #* | *Brand* | *Reusable* | *Custom-made* |
|---|---|---|---|---|---|
| US machine | Vevo MD Imaging System | 51475 | Fujifilm Sonosite, Inc. | Y | N |
| US transducers | UHF70 Transducer | 51416 | Fujifilm Sonosite, Inc. | Y | N |
| | UHF48 Transducer | 51415 | Fujifilm Sonosite, Inc. | | |
| | UHF22 Transducer | 51414 | Fujifilm Sonosite, Inc. | | |

***Other***

| *Product* | *Name* | *Part #* | *Brand* | *Reusable* | *Custom-made* |
|---|---|---|---|---|---|
| Isofluran | Multiple sources | n/a | n/a | N | N |

| | | | | | |
|---|---|---|---|---|---|
| Hair removal creme | Nair Body Cream | n/a | Church & Dwight Co., Inc. | N | N |
| Eye lubricant | Multiple sources | n/a | n/a | N | N |
| Ethanol wipes | Multiple sources | n/a | n/a | N | N |
| Povidone iodine | Betadine Surgical Scrub (7.5% povidone-iodine) Antiseptic Non-Sterile Scrub | 67618-154-16 | Purdue Products LP | N | N |
| Lidocaine | Multiple sources | n/a | n/a | N | N |
| 2x2 Gauze | Pivetal Non-Woven Gauze Sponge 2in. x 2in. (Gauze) | 21295051 | Patterson Veterinary | N | N |
| 4x4 Gauze | Pivetal Non-Woven Gauze Sponge 4in. x 4in. (Gauze) | 21295051 | Patterson Veterinary | N | N |
| Hair trimmer | Bravmini+ Purple | 41590-0438 | WAHL | Y | N |
| Hair trimmer blade | Bravmini+ Designer Blade | 41590-7840 | WAHL | Y | N |
| Tong depressors | Sterile Regular Tongue Depressors (Tongue Depressors) | 25-705 | Puritan Medical Products | N | N |
| Cotton Swab | Sterile Cotton Tipped Applicators (Sterile Applicators) | 56800 | AMD-Ritmed | N | N |
| Tape | Transpore Surgical Tape (tape) | 1527-1 | 3M | N | N |

| ***Experimental models: organisms/strains*** | **Strain** | ***Supplier*** |
|---|---|---|
| Rattus norvegicus | CD® IGS (Sprague Dawley) Rats | Charles River |
| Callithrix jacchus | | Worldwide Primates Inc. and in-house breeding colony |

**Supplementary Table 4: Histological processing**

| Reagent | Source | Identifier |
|---|---|---|
| ***Chemicals*** | | |
| Paraformaldehyde | Multiple sources | Cas# 30525-89-4 |
| Glycerol | Sigma | Cat# G5516, Cas# 56-81-5 |
| Tissue-Tek® O.C.T Compound | Electron Microscopy Science (Sakura Finetek) | Cat# 62550-12 |
| Fluoromount™ Aqueous Mounting Medium | Sigma | Cat# F4680 |
| 4',6-diamidino-2-phenylindole (DAPI) | Thermo Fisher Scientific | Cat# 62248 |
| Ethyl alcohol, Pure 200 proof. | Multiple brands | Cas# 64-17-5 |
| Sodium chloride | Multiple brands | Cas# 7647-14-5 |
| Sodium dodecyl sulfate | Sigma | Cat# 75746, Cas# 151-21-3 |
| Boric acid | Thermo Scientific Chemicals | Cat# 12680, Cas# 10043-35-3 |
| Sodium sulfite | Sigma-Aldrich | Cat# S0505, Cas# 7757-83-7 |
| Antipyrine Crystalline | Sigma-Aldrich | Cat# A5882, Cas# 60-80-0 |
| Nicotinamide | Sigma | Cat# 72340, Cas# 98-92-0 |
| N-butyldiethanolamine | Sigma | Cat# 471240, Cas# 102-79-4 |

| ***Primary antibodies*** | | ***Identifier*** | ***Working dilution*** |
|---|---|---|---|
| Chicken anti-Green Fluorescent Protein Antibody | Aves Labs | Cat# GFP-1020, RRID:AB_100002 | 1:5000 |
| Guinea pig anti-NeuN Antibody | Millipore | Cat# ABN90P, RRID:AB_2341095 | 1:5000 |
| Mouse anti-NeuN Antibody | Millipore | Cat# MAB377, RRID:AB_2298772 | 1:5000 |
| Mouse anti-parvalbumin Antibody | Swant | Cat# 235, RRID:AB_10000343 | 1:2000 |

| ***Secondary antibodies*** | ***Brand*** | ***Identifier*** | ***Working dilution*** |
|---|---|---|---|
| Goat anti-Guinea Pig IgG (H+L) Highly Cross-Adsorbed Secondary Antibody, Alexa Fluor™ 555 | Thermo Fisher Scientific | Cat# A-21435, RRID:AB_2535856 | 1:1000 |
| Goat anti-Chicken IgY (H+L) Secondary Antibody, Alexa Fluor 488 | Thermo Fisher Scientific | Cat# A-11039, RRID:AB_2534096 | 1:1000 |
| Goat anti-Mouse IgG (H+L) Highly Cross-Adsorbed Secondary Antibody, Alexa Fluor™ 647 | Thermo Fisher Scientific | Cat# A-21236, RRID:AB_2535805 | 1:1000 |
